# A kinase-dependent checkpoint prevents escape of immature ribosomes into the translating pool

**DOI:** 10.1101/656942

**Authors:** Melissa D. Parker, Jason C. Collins, Boguslawa Korona, Homa Ghalei, Katrin Karbstein

## Abstract

Premature release of nascent ribosomes into the translating pool must be prevented, as these do not support viability and may be prone to mistakes. Here we show that the kinase Rio1, the nuclease Nob1, and its binding partner Pno1 cooperate to establish a checkpoint that prevents the escape of immature ribosomes into polysomes. Nob1 blocks mRNA recruitment, and rRNA cleavage is required for its dissociation from nascent 40S subunits, thereby setting up a checkpoint for maturation. Rio1 releases Nob1 and Pno1 from pre-40S ribosomes to discharge nascent 40S into the translating pool. Weakly binding Nob1 and Pno1 mutants can bypass the requirement for Rio1, and Pno1 mutants rescue cell viability. In these strains, immature ribosomes escape into the translating pool, where they cause fidelity defects and perturb protein homeostasis. Thus, the Rio1-Nob1-Pno1 network establishes a checkpoint that safeguards against the release of immature ribosomes into the translating pool.

## Introduction

To maintain and balance protein levels within cells to support life, ribosomes must ensure that mRNA codons are faithfully translated into functional proteins. To guarantee their accurate function, the cell has to safeguard ribosome integrity during both assembly and its functional cycle. Ribosome assembly is a highly regulated process, involving the proper folding and processing of 4 rRNAs, as well as binding of 79 ribosomal proteins (r-proteins). Assembly is facilitated by over 200 transiently binding assembly factors that promote assembly and quality control, and prevent immature ribosomes from initiating translation prematurely (Kressler et al., 2017; Pena et al., 2017; Warner, 1999; Woolford and Baserga, 2013).

To prevent misassembled ribosomes from reaching the translating pool, the premature small (pre-40S) ribosomal subunit undergoes a series of quality control checkpoints during late cytoplasmic maturation that verify proper ribosomal structure and function (Collins et al., 2018; Ghalei et al., 2017; Strunk et al., 2012). The importance of these checkpoints for cellular function is illustrated by the numerous diseases caused by haploinsufficiency or mutations in ribosomal proteins and assembly factors. These alterations dysregulate ribosome concentrations, and/or lead to misassembled ribosomes and an increased propensity of patients to develop cancer (Bolze et al., 2013; Burwick et al., 2011; Ellis and Lipton, 2008; Freed et al., 2010; Goudarzi and Lindstrom, 2016; Vlachos et al., 2012).

One of the final steps in the biogenesis of 40S subunits in yeast is the maturation of the 3’-end of 18S rRNA from its precursor, 20S pre-rRNA. This step is carried out by the essential endonuclease Nob1 (Fatica et al., 2003; Fatica et al., 2004; Lamanna and Karbstein, 2009; Pertschy et al., 2009) and is promoted by its direct binding partner Pno1 (Woolls et al., 2011). Pno1 also blocks the premature incorporation of Rps26, as these two proteins occupy the same location on nascent or mature ribosomes, respectively (Ameismeier et al., 2018; Heuer et al., 2017; Johnson et al., 2017; Scaiola et al., 2018; Strunk et al., 2011).

Rio1 is an essential aspartate kinase bound to very late cytoplasmic pre-40S subunits that have shed all bound assembly factors except Nob1 and Pno1 (Ferreira-Cerca et al., 2014; Hector et al., 2014; Turowski et al., 2014; Vanrobays et al., 2003; Widmann et al., 2012). Depletion of Rio1 or overexpression of a catalytically-inactive Rio1 mutant leads to the accumulation of 20S pre-rRNA and assembly factors in 80S-like ribosomes (Belhabich-Baumas et al., 2017; Ferreira-Cerca et al., 2014; Vanrobays et al., 2001; Widmann et al., 2012). However, the role Rio1 plays in 18S rRNA maturation and ribosome assembly remains unknown, despite its interest as a target for the development of anti-cancer drugs (Kiburu and LaRonde-LeBlanc, 2012; Kubinski et al., 2017; Mielecki et al., 2013; Read et al., 2013; Weinberg et al., 2017), and the observation that RIOK1 mutations accumulate in human cancers (TCGA Research Network: https://www.cancer.gov/tcga).

In this study, we use a combination of biochemical and genetic experiments to dissect the role of Rio1 in ribosome assembly. Our data show that Nob1 blocks the premature entry of nascent 40S subunits into the translating pool and requires rRNA maturation for its dissociation from nascent 40S subunits, thereby ensuring that only fully matured subunits engage in translation. Additionally, we provide evidence that Rio1 releases Nob1 and Pno1 from nascent ribosomes in an ATPase-dependent manner, and that weakly binding Nob1 and Pno1 mutants can bypass the requirement for Rio1. Thus, the Rio1 kinase and Nob1 nuclease cooperate to restrict and regulate the entry of nascent ribosomes into the translating pool only after they are properly matured. Finally, bypassing Rio1 via self-releasing mutations in Pno1 results in release of immature ribosomes containing pre-rRNA, but lacking Rps26 into the translating pool, where they produce fidelity defects and alter mRNA specificity in yeast (Ferretti et al., 2017), but rescue the lethality of Rio1 depletion. Together, these data reveal the function of a disease-associated kinase in licensing the entry of only mature ribosomes into the translating pool, thereby safeguarding the integrity of translating ribosomes.

## Results

### Nob1 inhibits mRNA recruitment

Nob1 is the endonuclease responsible for the final cleavage of pre-18S (20S) rRNA to produce its mature 3’-end (Fatica et al., 2003; Fatica et al., 2004; Lamanna and Karbstein, 2009). Thus, in Nob1-depleted cells, ribosomes containing 20S pre-rRNA accumulate (Fatica et al., 2003; Fatica et al., 2004; Lamanna and Karbstein, 2009) and are found in polysomes (**Figure 1A**) (Soudet et al., 2010; Strunk et al., 2012). This is surprising because Nob1 is an essential gene (**Figure S1A**). To test if the discrepancy between active translation and lack of viability arises because the gene products from immature ribosomes poison the cell, we used a dominant-negative catalytically-inactive mutant of Nob1 (Nob1-D15N) (**Figure 1B**). Nob1-D15N is a mutation in the conserved PIN domain of Nob1, rendering Nob1 able to bind but not cleave its 20S pre-rRNA substrate (Fatica et al., 2004; Pertschy et al., 2009). 20S pre-RNA accumulation in wild type cells is noticeable after Gal-driven overexpression of Nob1-D15N for 8 hours, but not in cells expressing an empty vector (**Figure 1C**).

**Figure 1.**
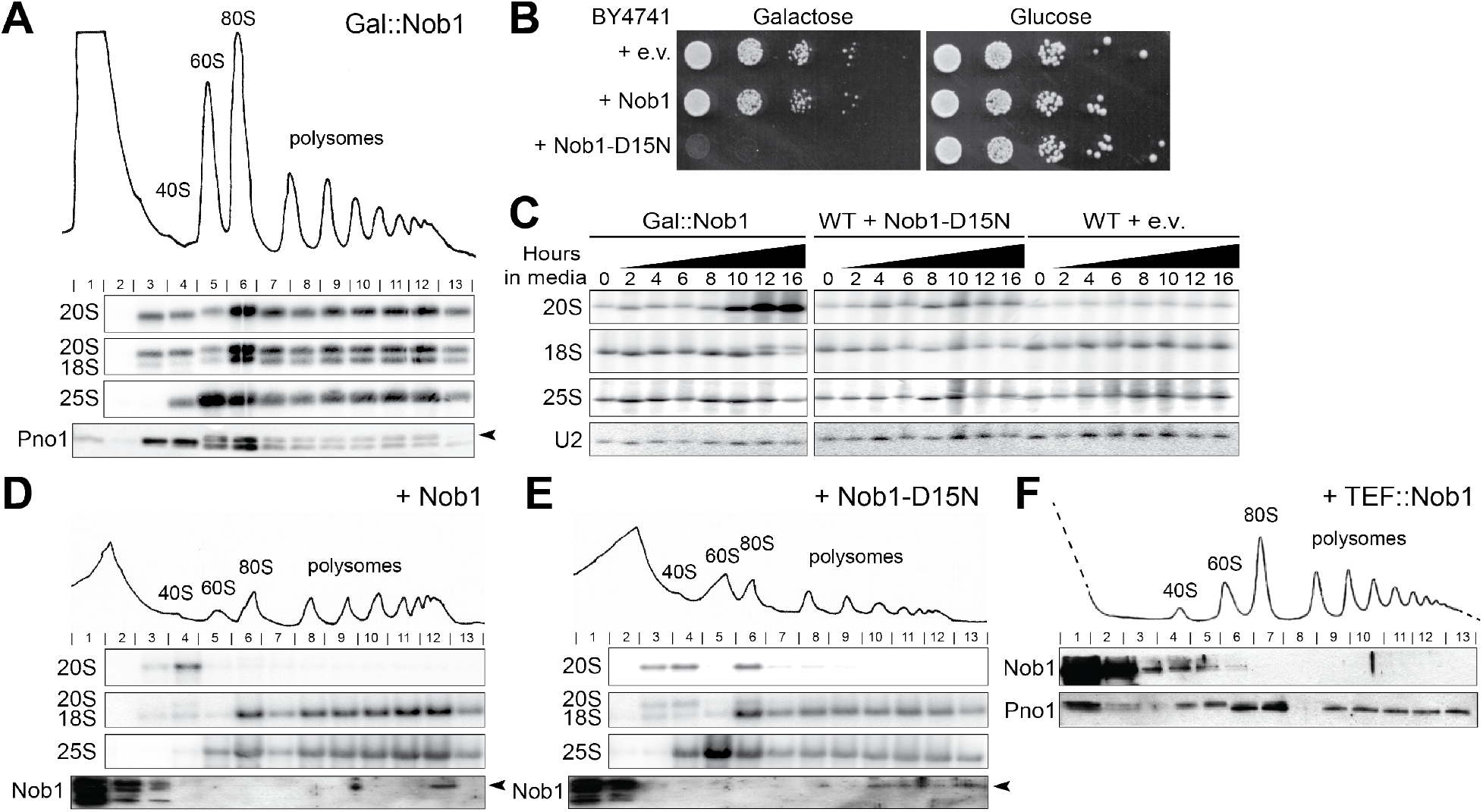
Nob1 prevents entry of pre-40S subunits into the polysomes. **A)** 10%–50% sucrose gradients from lysates of cells depleted of Nob1 by growth in glucose for 12 hr. Shown below the absorbance profile at 254 nm are Northern blots of pre-18S rRNA (20S), mature 18S, and 25S rRNAs and Western blots probing for Pno1. Arrowhead notes the upper band corresponding to Pno1. **B)** Growth of wild type yeast cells transformed with an empty vector (e.v.) or Nob1 or Nob1-D15N under the galactose-inducible, glucose-repressible Gal promoter were compared by 10-fold serial dilutions on glucose or galactose dropout plates. **C)** Northern blot analyses of total cellular RNA from cells depleted of Nob1 grown in glucose for the indicated times, and total cellular RNA from wild type (WT) BY4741 cells overexpressing Nob1-D15N or transformed with an empty vector grown in galactose for the indicated times. **D, E)** Sucrose gradients of wild type BY4741 cells overexpressing Nob1 (D) or Nob1-D15N (E) in galactose for 12 hr. Western blots probed for Nob1. Arrowhead notes the upper band corresponding to Nob1. **F)** Sucrose gradients of wild type BY4741 cells overexpressing Nob1 under a Tef2 promoter grown in glucose. See also **Figure S1**.

To assess whether the Nob1-containing pre-40S ribosomes entered the polysomes, as observed for pre-ribosomes accumulated in the absence of Nob1, we performed polysome profiling followed by Northern blotting on cells overexpressing Nob1 or Nob1-D15N. Overexpressing wild-type Nob1 results in 20S pre-rRNA concentrated only in the 40S fraction, while the polysomes contained only mature 18S rRNA (**Figure 1D**). In contrast to Nob1-depleted cells, very little 20S pre-rRNA escaped into the polysomes in Nob1-D15N-overexpressing cells, and instead accumulated in pre-40S and 80S-like ribosome peaks (**Figure 1E**). This observation is also consistent with the appearance of robust polysomes in Nob1-depleted cells, but not Nob1-D15N cells. Thus, ribosomes containing immature 20S pre-rRNA can recruit mRNA to enter the polysomes in the absence of Nob1, but not its presence, suggesting that Nob1 blocks mRNA recruitment.

If Nob1 blocks mRNA recruitment, then it should not be found in the polysomes. Consistently, Nob1 is not found in the polysomes of wild type cells (Strunk et al., 2012), or when expressed under the Cyc1 and Tef2 promoters, which produce slightly less and significantly more Nob1 than the endogenous promoter, respectively (**Figure 1F and 4E**).

We also considered the possibility that it is not the presence of Nob1, but rather its interacting partner Pno1 that blocks entry into the polysomes. However, we note that Pno1 can be found in the polysomes in Nob1-depleted cells, showing that Pno1 does not block polysome recruitment (**Figure 1A**). The finding that Pno1 remains bound to actively translating 20S-containing ribosomes in Nob1-depleted cells also explains why these translating ribosomes do not support growth. Pno1 blocks Rps26 binding (Heuer et al., 2017; Strunk et al., 2011), and thus, the remaining Pno1 will prevent binding of Rps26, an essential protein required for translation of ribosome components (Ferretti et al., 2017).

### Nob1 release requires rRNA cleavage

If Nob1 release from nascent 40S requires rRNA cleavage by Nob1, then Nob1 blocking mRNA recruitment to premature ribosomes would enable a quality control mechanism to ensure only ribosomes containing matured rRNA enter the polysomes. To test whether Nob1-dependent rRNA cleavage is required for its dissociation from ribosomes, we used a previously described quantitative *in vitro* RNA binding assay (Lamanna and Karbstein, 2009). This assay uses native gel-shift to measure the binding of Nob1 to mimics of the 20S pre-rRNA substrate (H44-A2), the 18S rRNA ribosome product (H44-D), or the 3’-ITS1 product (D-A2). The data show that Nob1 binds the substrate mimic and the 3’-ITS1 product with similar affinities (*K*_d_= 0.93 and 0.96 μM, respectively). In contrast, Nob1 binds the 18S rRNA mimic more weakly (*K*_d_ =1.89 μM) (**Figure 2**). This finding shows that Nob1 predominantly interacts with ITS1, consistent with previous structure probing data (Lamanna and Karbstein, 2009; Turowski et al., 2014). Furthermore, the data suggest that after Nob1 cleavages 20S pre-RNAs, Nob1 will remain bound to its 3’-cleavage product and not to the matured 18S rRNA. Thus, Nob1 blocks premature ribosomes from binding mRNA until rRNA is cleaved, allowing Nob1 to dissociate from nascent ribosomes, and thereby setting up a mechanism to ensure only ribosomes with fully matured rRNA enter the translating pool.

**Figure 2.**
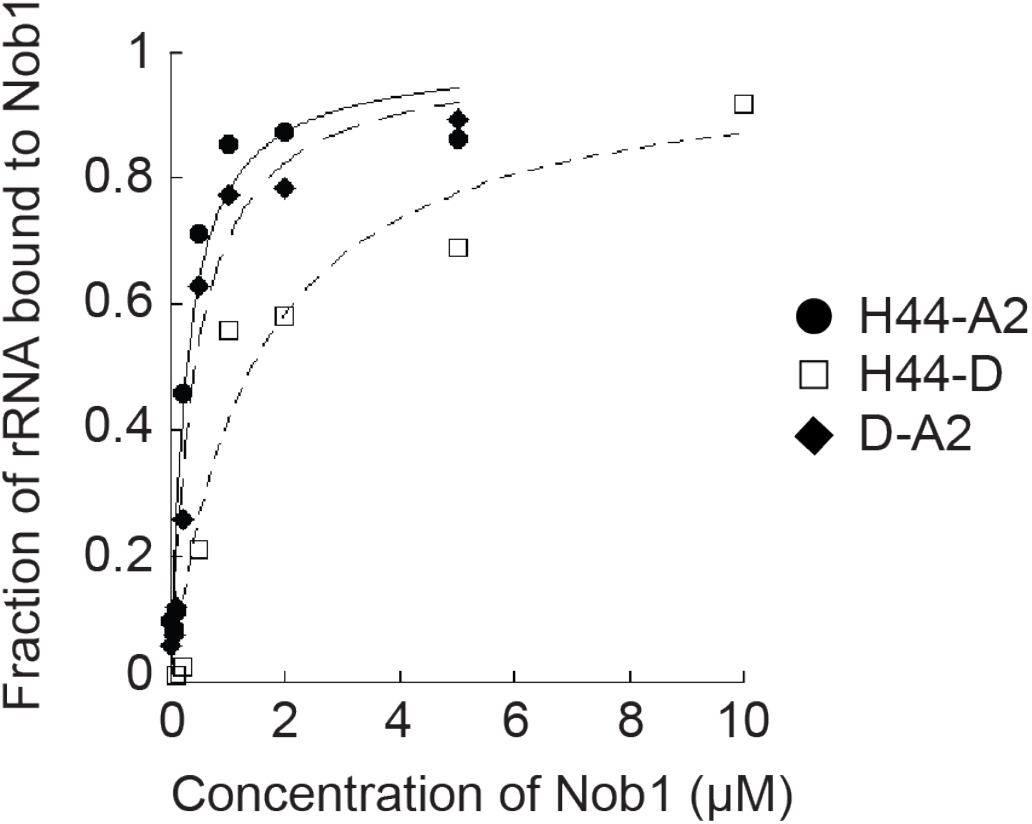
Nob1 dissociates with the 3’-cleavage product following rRNA cleavage. Representative RNA binding assay with *in vitro* transcribed H44-A2 (20S pre-rRNA mimic, black circles), H44-D (18S rRNA mimic, white squares), or D-A2 (3’-ITS1, black diamonds) RNAs and with recombinant Nob1. Four or five independent experiments yielded values of *K*_d_ = 0.93 +/− 0.09 μM for Nob1 binding H44-A2, *K*_d_ = 0.96 +/− 0.05 μM for Nob1 binding D-A2, and *K*_d_ = 1.89 +/− 1.04 μM for Nob1 binding H44-D.

### Rio1 authorizes translation initiation of nascent 40S ribosomes

Nevertheless, because Nob1 is also bound to Pno1 (Ameismeier et al., 2018; Woolls et al., 2011), Nob1 release from pre-40S also requires separation from Pno1. Thus, to test if other late-acting 40S assembly factors play a direct role in Nob1 release, we carried out a limited screen for factors whose overexpression rescues the dominant-negative phenotype from Nob1-D15N overexpression (data not shown). This screen showed that overexpression of the aspartate kinase Rio1 rescues the growth phenotype from Nob1-D15N overexpression (**Figure 3A**). Furthermore, Rio1 activity is required for this rescue, as mutations that block phosphorylation, D261A (the phosphoaspartate) (Ferreira-Cerca et al., 2014) and K86A (in the P-loop), did not rescue the Nob1-D15N growth phenotype (**Figure 3A**).

**Figure 3.**
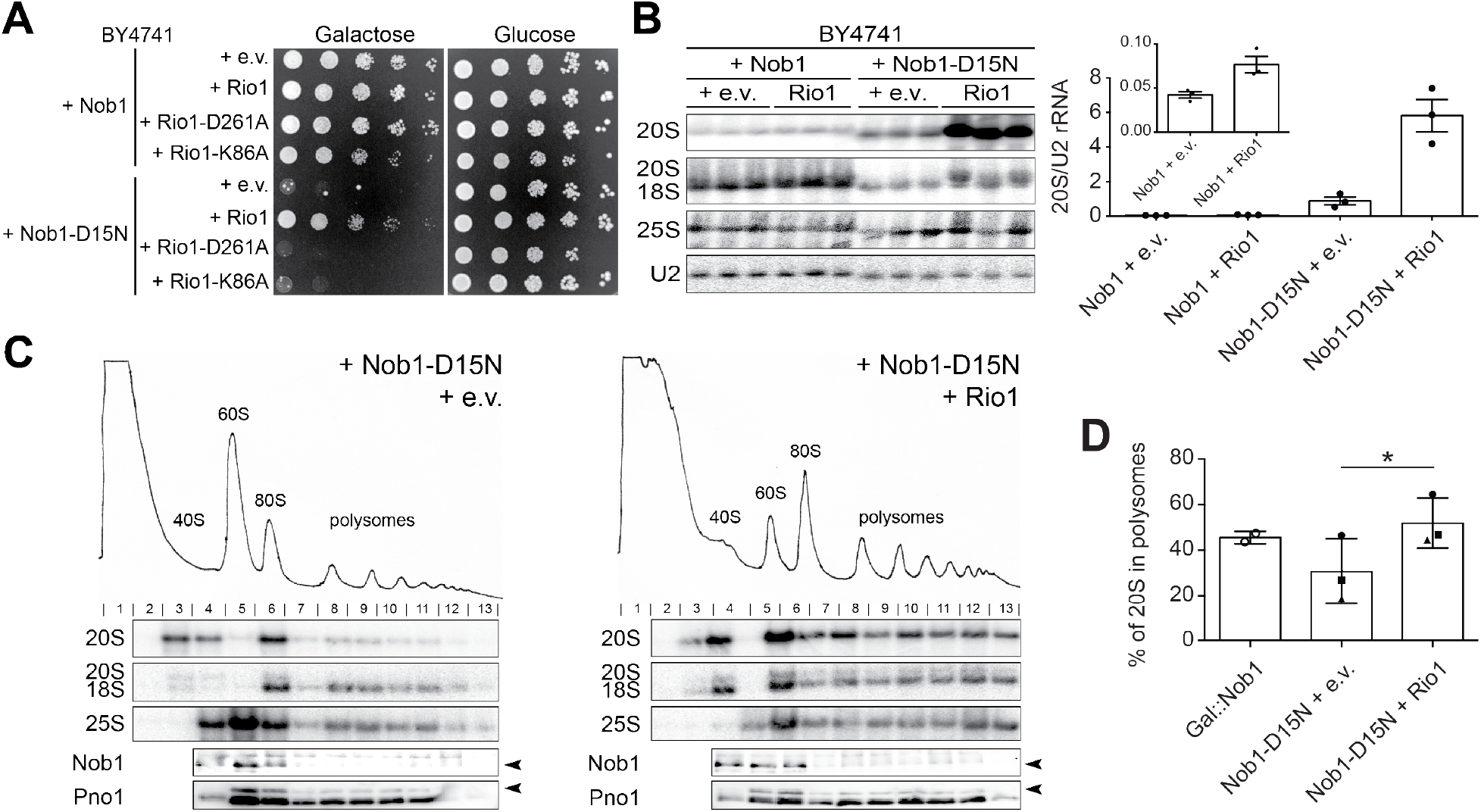
Rio1 releases 40S ribosomes into the translating pool. **A)** Growth of cells containing an empty vector or wild-type Rio1, Rio1-D261A, or Rio1-K86A under a copper-inducible (Cup1) promoter, and Nob1 or Nob1-D15N under the Gal promoter were compared by 10-fold serial dilutions on glucose or galactose dropout plates with 100 μM CuSO_4_. **B)** Left: Northern blot analyses of total cellular RNA from cells in panel A grown in galactose with 100 μM CuSO_4_ for 16 hr. Right: Quantification of the Northern blot. 20S pre-rRNA levels were normalized to U2. **C)** 10%-50% sucrose gradients from cell lysates of cells in panel B. Northern blots of 20S, 18S, and 25S rRNA and Western blots probing for Nob1 and Pno1 are shown below the absorbance profile at 254 nm. Arrowheads note the bands corresponding to Nob1 (lower band) and Pno1 (upper band). **D)** Quantification of the gradient Northern blots in C and in Figure 1A. Percentage of 20S pre-rRNA in polysomes (fractions 8-13) compared to total 20S pre-rRNA was calculated. Data are the average of three biological replicates, and error bars indicate SEM. Samples grown and analyzed on the same day were considered paired replicates, as indicated by the circle, square, and triangle dots on the graph. Paired t-test was used for statistical analysis; *p=0.0123. See also **Figure S1**.

To test if Rio1 overexpression promotes endonuclease activity of Nob1 and thus rescues the Nob1 mutation by “repairing” its catalytic activity, we carried out Northern analysis. Overexpressing Nob1-D15N and Rio1 together resulted in a 6.5 fold increase in 20S pre-rRNA accumulation compared to Nob1-D15N alone (**Figure 3B**). This is the opposite of what would be expected if Rio1 “repairs” Nob1 activity. Additionally, overexpression of Rio1 did not rescue the lack of Nob1 (**Figure S1A**), as expected if Rio1’s role is to release Nob1 rather than to promote rRNA cleavage. These data show that Rio1 does not rescue the growth phenotype of Nob1-D15N by stimulating rRNA cleavage.

To test if instead Rio1 overexpression rescues the Nob1-D15N growth phenotype by releasing Nob1, thereby allowing 20S pre-rRNA-containing ribosomes to enter the translating pool as in Nob1-depleted cells, we used polysome profiling coupled with Northern analysis. As before, Nob1-D15N overexpression results in accumulation of 20S pre-rRNA in pre-40S and 80S-like ribosomes (**Figure 3C**, left), with only 30% of 20S pre-rRNA in polysome fractions (**Figure 3D**). Simultaneous Rio1 overexpression releases 20S pre-rRNA-containing ribosomes into the polysomes (**Figure 3C**, right), with a statistically significant increase to 52% of 20S pre-rRNA in polysomes (**Figure 3D**). The accumulation of pre-ribosomes in the translating pool when Rio1 is overexpressed in the Nob1-D15N background is the same as that observed upon Nob1 depletion (52% and 46%, respectively). These data show that Rio1 overexpression promotes the release of immature, 20S-containing ribosomes into the translating pool. Furthermore, the data suggest that the mechanism by which this occurs is via Nob1 release, thereby turning Nob1-D15N-containing ribosomes into Nob1-depleted ribosomes. This model is further supported by polysome analysis of Rio1 depleted cells, where only 19% of 20S pre-rRNA-containing ribosomes reach the polysomes, and these mRNA-bound ribosomes lack Nob1 (**Figure S1B**).

### Rio1 binds Nob1 and Pno1 directly and stimulates their release from pre-40S ribosomes

Rio1 is an atypical aspartate kinase. By analogy to its close relative, Rio2, it is believed that Rio1 phosphorylation functions as a switch, which is reset after hydrolysis (Ferreira-Cerca et al., 2012; Knuppel et al., 2017). Previous analysis suggests that Rio1 interacts with pre-40S ribosomes during the final cytoplasmic assembly steps when the pre-40S is bound only to Nob1 and its binding partner Pno1 (Turowski et al., 2014; Widmann et al., 2012), consistent with our data that indicate a role for Rio1 in Nob1 release to allow for discharge of the nascent 40S subunits into the translating pool.

To test this model, we performed *in vitro* protein binding assays with recombinant Rio1, Nob1, and Pno1. These experiments show that MBP-Rio1 binds Nob1 and Pno1, but not either Nob1 or Pno1 individually, suggesting that Rio1 recognizes the Nob1-Pno1 complex (**Figure 4A** and **S2**). Importantly, the presence of AMPPNP, a non-hydrolyzable ATP analog, is required for formation of the Rio1•Nob1•Pno1 complex, as little to no complex formation is observed in the presence of ADP (**Figure 4A** and **S2**).

**Figure 4.**
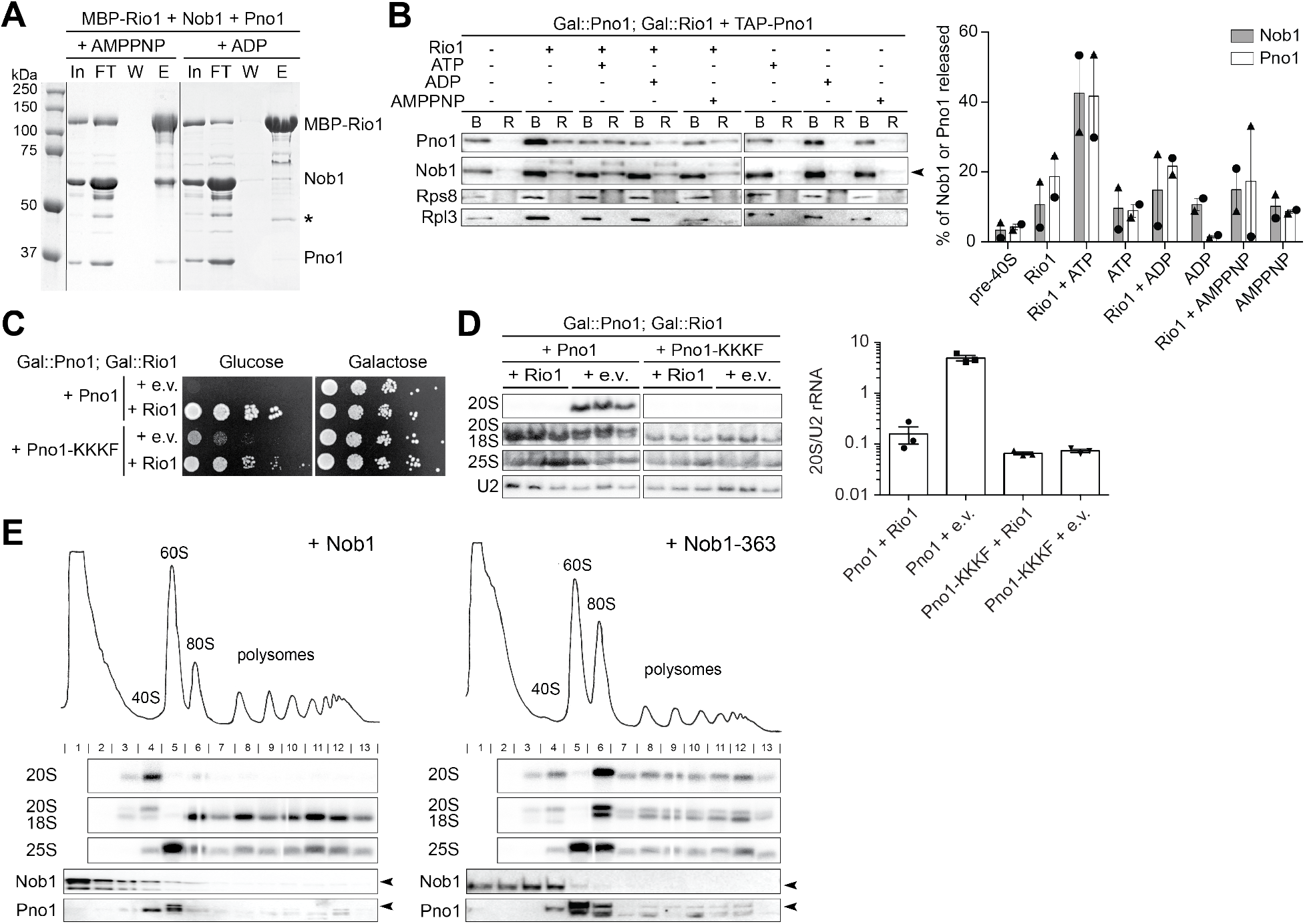
Rio1 stimulates release of Nob1 and Pno1 from the pre-40S ribosome. **A)** Rio1 binds Nob1 and Pno1 in the presence of the non-hydrolyzable ATP analog AMPPNP. Shown are Coomassie-stained SDS-PAGE gels of protein binding assays using purified, recombinant MBP-Rio1, Nob1, and Pno1 in the presence of AMPPNP or ADP. In, input; FT, flow-through; W, final wash; E, elution. The order of the samples was edited for clarity. *MBP. See also **Figure S2. B)** Left: Western blot analysis of a release assay measuring ribosome-bound (B) proteins and released (R) proteins upon the addition of purified, recombinant Rio1 and ATP, ADP, or AMPPNP. Right: Quantification of the release assay Western blot. The percent of Nob1 and Pno1 released from the ribosome compared to the total Nob1 or Pno1 in each sample. Data are the average of two biological replicates, as indicated by the circle and triangle dots on the graph, and error bars indicate SEM. **C)** Growth of cells expressing either wild-type Pno1 or Pno1-KKKF (K208E/K211E/K213E/F214A) and either an empty vector or Rio1 were compared by 10-fold serial dilutions on glucose or galactose dropout plates. See also **Figure S3. D)** Left: Northern blot analyses of total cellular RNA from cells in panel C grown in glucose for 16hr. Samples were run on the same gel and the order was edited for clarity. Right: 20S pre-rRNA accumulation was normalized to U2 levels in these cells. **E)** Sucrose gradients analyzed by Northern and Western blot of lysate from Gal::Nob1; Gal::Rio1 cells expressing wild type Nob1 (left) or the Nob1-363 truncated protein (right) on a plasmid. Arrowheads note the bands corresponding to Nob1 and Pno1. See also **Figure S4** and **Figure S6**.

These data suggest Rio1 recognizes and binds the interface of the Nob1•Pno1 complex in an ATP-dependent manner. To test if autophosphorylation (and therefore ATP hydrolysis) is responsible for breaking this complex, we developed an *in vitro* release assay using assembly intermediates purified from yeast and purified recombinant Rio1. In this assay, TAP-Pno1 ribosomes purified from cells depleted of Rio1 are incubated with Rio1 in the presence of ATP, AMPPNP, or ADP. Release of assembly factors was monitored using an assay where the reactions are layered onto a sucrose cushion to pellet ribosomes and all bound factors, while free proteins will be in the supernatant. Little Nob1 or Pno1 (2% and 5% of Nob1 or Pno1, respectively) were released in a mock incubation (**Figure 4B**). Addition of Rio1 alone, or in the presence of ADP or AMPPNP increased this slightly (~10% of Nob1 and 20% Pno1 released, respectively), while addition of Rio1 and ATP led to a >10-fold increase in the release of these assembly factors (32% Nob1 and 54% Pno1, respectively, **Figure 4B**). This finding demonstrates that Rio1 uses ATP hydrolysis to stimulate the dissociation of Nob1 and Pno1 from the pre-40S subunit.

### Weakly binding Nob1 and Pno1 mutants can bypass Rio1 activity

The data above show that Rio1 can release Nob1 and Pno1 from nascent 40S subunits *in vitro*. To confirm a role for Rio1 in the release of Nob1 and Pno1 from pre-40S ribosomes *in vivo*, we screened a collection of mutants in Pno1 and Nob1 for their ability to rescue the loss of cell viability upon Rio1 depletion. These included Pno1 mutants that disrupt the binding to Nob1 (GXXG, WK/A, HR/E, DDD/K) (Woolls et al., 2011) (**Figure S3A**), or weakened its interactions with ribosome (KKKF) (Johnson et al., 2017) (**Figure 4C**), as well as mutations in Nob1 that weaken ribosome binding (truncations Nob1-434 and Nob1-363, and L88A/S89A, L93A/L96A, K320A/F322A, Q28R, D271N, D271R/F272A, Q280R, T225A, K80A, Y300A, and R303A), or Pno1 binding (W223G) (Sturm et al., 2017) (**Figure S3B** and **S4A**). Finally, overexpression of Nob1 was also tested (**Figure S3C**). Of all of these mutants, only the weak-binding Pno1-KKKF (K208E/K211E/K213E/F214A) mutant was able to rescue the lethal phenotype from Rio1 depletion (**Figure 4C, S3**, and **S4A**).

To confirm that Pno1-KKKF rescued Rio1 depletion, we analyzed pre-rRNA levels in cells expressing wild type (wt) Pno1 or Pno1-KKKF in the presence or absence of Rio1 using Northern blotting. These data showed a 30-fold increase in 20S pre-rRNA accumulation in cells lacking Rio1 compared to cells expressing wt Rio1 and wt Pno1. In contrast, with Pno1-KKKF, no 20S pre-rRNA accumulation was observed in the absence of Rio1 (**Figure 4D**). Together these data show that the function of Rio1 can be bypassed by the release of Pno1 from the pre-40S ribosome.

Surprisingly, while none of the self-releasing Nob1 mutants rescued the growth defects observed from Rio1 depletion, this could be explained if Pno1 remained bound to ribosomes in the polysomes of these strains, as it does in Gal::Nob1 cells (**Figure 1A**), thereby blocking Rps26 recruitment (Heuer et al., 2017; Strunk et al., 2011). To test this idea, we analyzed polysome profiles from cells depleted of Rio1 while expressing either full length Nob1 or a truncated Nob1-363 (in which the Nob1 gene does not encode for amino acids 364-459). Nob1-363 binds rRNA more weakly as suggested by a growth phenotype from this mutant, which can be rescued when Nob1 is overexpressed from the TEF promoter (**Figure S4A**), as well as RNA binding data (**Figure S4B**). While 20S pre-rRNA accumulates in the 40S and 80S fractions in cells depleted of Rio1 but containing wild type Nob1, both 20S pre-rRNA and Pno1 escape into the polysomes when Nob1-363 is expressed (**Figure 4E**). Thus, this weakly-binding, truncated Nob1 mutant can bypass the requirement of Rio1 for the dissociation of Nob1 from nascent 40S, providing strong support for a role for Rio1 in Nob1 release.

### Bypass of Rio1 activity leads to release of immature 40S ribosomes into the translating pool

The data above provide strong evidence for a role for the kinase Rio1 in releasing Nob1 and Pno1 from nascent 40S subunits. Because Nob1 dissociation also requires prior Nob1-dependent rRNA cleavage, this pathway ensures only ribosomes containing fully matured rRNA are discharged into the translating pool. Thus, these data strongly support a role for Rio1 in ensuring only matured ribosomes enter the translating pool. To test this directly, we took advantage of the self-releasing Pno1-KKKF mutant, which bypasses the requirement for Rio1 and allows for cellular growth in the absence of Rio1. If Rio1 restricts premature entry of immature ribosomes into the translating pool, we would predict that bypassing Rio1 with the Pno1-KKKF mutant would allow for the escape of immature ribosomes into the polysomes.

To test this prediction, we analyzed the rRNA in polysomes of cells expressing wild type Pno1, or Pno1-KKKF. In cells expressing Pno1-KKKF, 15% of 20S pre-rRNA escaped into the polysomes compared to only 3% of 20S pre-rRNA in wild type cells (**Figure 5A** and **5B**). Importantly, this is not because there is accumulation of 20S pre-rRNA in the Pno1-KKKF mutant (**Figure 4D**). This finding confirms a role for Rio1 in restricting the release of immature ribosomes into the translating pool.

**Figure 5.**
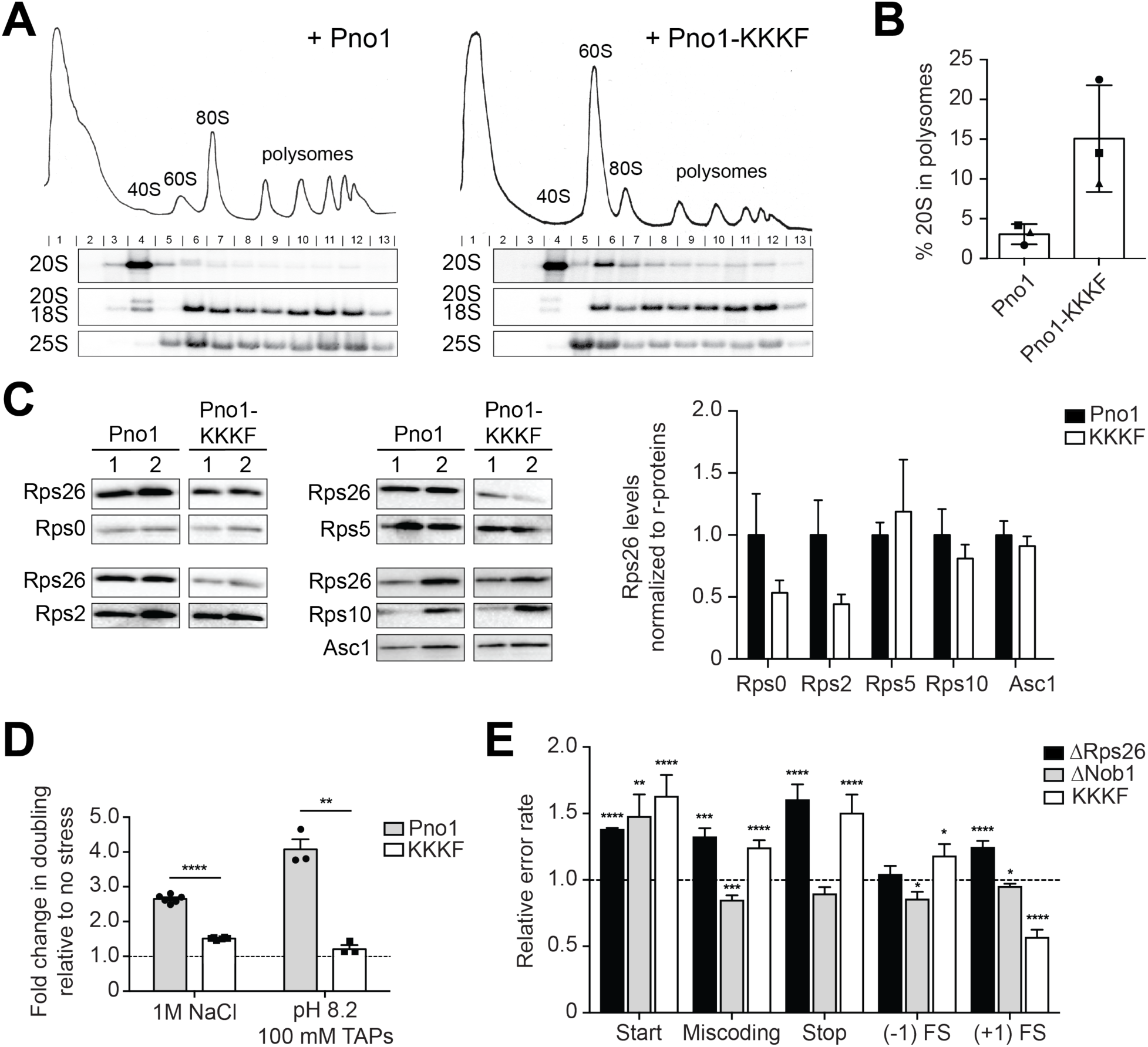
Rio1-mediated quality control ensures only mature 40S are released into the translating pool. **A)** 10-50% sucrose gradients of cell lysates from Gal::Rio1; Gal::Pno1 cells expressing wt Rio1 and either wt Pno1 or Pno1-KKKF from plasmids. Shown below the absorbance profile at 254 nm are Northern blots of 20S, 18S, and 25S rRNAs and Western blots probing for Nob1 and Pno1. Arrowheads note the bands corresponding to Nob1 and Pno1. **B)** Quantification of the gradient Northern blots. The percentage of 20S pre-rRNA in polysomes (fractions 8-13) compared to total 20S pre-rRNA was calculated. Data are the average of three biological replicates, as indicated by the circle, square, and triangle dots on the graph, and error bars indicate SEM. **C)** Left: Western blot analyses of ribosomal proteins from ribosomes purified from cells expressing wild-type Pno1 or Pno1-KKKF. Right: Quantification of Western blots of 2-7 technical replicates for each of 2 biological replicates. Rps26 levels were normalized to the indicated ribosomal protein levels. Error bars indicate SEM. **D)** Changes in doubling time in cells expressing Pno1 or Pno1-KKKF after exposure to high salt or high pH. Values were compared to no-stress conditions (fold change = 1). Data are the average of three (high pH) or six (high salt) biological replicates and error bars indicate SEM. **p < 0.005, ****p < 0.0001 by unpaired t-test. Compare to the doubling time of cells deficient in Rps26 after exposed to high salt or high pH (Ferretti et al., 2017). **E)** The effects from the Pno1-KKKF mutation, depletion of Rps26 (ΔRps26), and depletion of Nob1 (ΔNob1) on the fidelity of start codon recognition, decoding, stop codon recognition, and frameshifting (−1 and +1) were assayed using dual luciferase reporters. Shown are the relative error rates of the mutant (or depleted) samples relative to wild-type (replete) samples. Rps26 was depleted by growth in 100 ng/mL Dox (Ferretti et al., 2017) and Nob1 was depleted in glucose. Data are the average of 5-27 biological replicates, and error bars indicate the SEM. *p < 0.05, **p < 0.01, ***p < 0.001, and ****p < 0.0001 by unpaired t-test. See also **Figure S5**.

### Rio1 regulates Rps26 incorporation to form the mature ribosome

Premature binding of Rps26 is blocked by Pno1. Furthermore, binding of Rps26 is also incompatible with the path of the pre-rRNA (**Figure S5**), suggesting that 20S-containing ribosomes might lack Rps26, even if they do not contain Pno1. To test this notion, we probed Rps26 levels in ribosomes purified from cells expressing wt Pno1 or Pno1-KKKF. As shown above, Pno1-KKKF self-releases from these ribosomes, leading to the release of immature ribosomes containing 20S pre-rRNA, but lacking Pno1, into the polysomes. Western blot analysis relative to five other ribosomal proteins, Rps0, Rps2, Rps5, Rps10, and Asc1, showed that the Rps26 occupancy was reduced in ribosomes from cells expressing Pno1-KKKF compared to ribosomes from cells expressing wild type Pno1 (**Figure 5C**). This finding suggests that ribosomes containing immature 20S pre-rRNA lack Rps26, indicating that rRNA maturation is required for Rps26-incorporation.

To further test if the immature ribosomes in Pno1-KKKF cells lack Rps26 we took advantage of the previously characterized phenotypes of Rps26-deficiency (Ferretti et al., 2017). Rps26-deficient yeast cells accumulate Rps26-deficient ribosomes, which enable preferential translation of mRNAs with -4G residues in the Kozak sequence, including components of the Hog1 MAP-kinase pathway and the Rim101 pathway, which respond to high salt and high pH stress (Ferretti et al., 2017). As a result, accumulation of Rps26-deficient ribosomes leads to high salt and pH resistance (Ferretti et al., 2018; Ferretti et al., 2017). We therefore tested if cells expressing Pno1-KKKF are more resistant to high salt and high pH stress than their wild-type counterparts, as predicted if they release Rps26-deficient ribosomes into the translating pool. Indeed, similar to cells depleted of Rps26 (Ferretti et al., 2017), cells expressing Pno1-KKKF are resistant to high salt and high pH stress (**Figure 5D**). Together, these data demonstrate that bypass of Rio1 using the self-releasing Pno1-KKKF mutant allows ribosomes lacking Rps26 and containing immature rRNA to escape into the translating pool. Furthermore, the data also strongly suggest that Pno1-release *and* Nob1-dependent maturation of pre-rRNA are both required for Rps26-incorporation.

### Rio1 bypass limits translational fidelity

The data above demonstrate that bypass of Rio1 allows Rps26-deficient, 20S pre-rRNA-containing immature ribosomes to escape into the translating pool. To test if these ribosomes cause defects in translational fidelity, we took advantage of a collection of previously described luciferase reporter plasmids (Cheung et al., 2007; Harger and Dinman, 2003; Keeling et al., 2004; Salas-Marco and Bedwell, 2005). For these plasmids, firefly luciferase production depends on a mistranslation event. To decipher effects on translational fidelity that arise from the Pno1-KKKF mutant, we compared mistranslation frequency in the Pno1-KKKF background relative to wt Pno1. The Pno1-KKKF mutation produced defects on codon-selection, largely mirroring the defects obtained from Rps26-deficient cells (**Figure 5E**). In addition, the defects in start codon recognition and decreased +1 frameshifting produced by the Pno1-KKKF mutation resemble the defects from Nob1-deficient cells (**Figure 5E**). Thus, because Rio1 releases immature, Rps26-deficient ribosomes into the translating pool, while Rps26 or Nob1 depletion releases only Rps26-deficient or immature ribosomes, respectively, the effects from Rio1 bypass on translation fidelity are a combination of the effects from Nob1 and Rps26 depletion.

## Discussion

### Nob1 blocks mRNA recruitment to 20S pre-rRNA-containing ribosomes

Nascent 40S subunits arrive in the cytoplasm bound to seven assembly factors, which block premature translation initiation on immature assembly intermediates by preventing the association of translation initiation factors (Strunk et al., 2012). These assembly factors are then released in a series of regulated steps that form part of a translation-like cycle, which couples the release of these factors to quality control steps (Collins et al., 2018; Ghalei et al., 2017; Strunk et al., 2012). Furthermore, when pre-mature ribosomes do escape into the translating pool, they are unable to support cell viability (Soudet et al., 2010; Strunk et al., 2012). Together these observations demonstrate the importance of preventing premature translation initiation by immature ribosomes. The data herein demonstrate that the discharge of ribosomes into the translating pool is a regulated quality control step during maturation of the small ribosomal subunit.

Nob1 is the endonuclease that produces mature 18S rRNA (Fatica et al., 2003; Lamanna and Karbstein, 2009). Thus, yeast lacking Nob1, or yeast overexpressing a dominant-negative, inactive Nob1, Nob1-D15N, both accumulate pre-18S rRNA (20S pre-rRNA) (Pertschy et al., 2009). Nonetheless, the data show that Nob1-depleted, 20S pre-rRNA-containing ribosomes enter the translating pool, while ribosomes bound to Nob1-D15N do not bind mRNA. This finding strongly suggests that mRNA recruitment by the nascent 40S subunit requires the prior dissociation of Nob1.

Nob1-mediated rRNA cleavage produces two products, the 18S-containing 40S subunit, and the ITS1 product, which is subsequently degraded by the exonuclease Xrn1 (Stevens et al., 1991). Our data demonstrate that the Nob1 binding affinities for the precursor rRNA and ITS1 product mimics are indistinguishable, and much stronger than for the mature 18S rRNA product. Thus, after cleavage, Nob1 is expected to remain bound to the ITS1 product and not the ribosome product, consistent with previous structure probing and cross-linking analyses (Lamanna and Karbstein, 2009; Turowski et al., 2014). Collectively, these findings support a model by which Nob1’s cleavage at the 3’-end of 18S rRNA is required for its dissociation from the nascent subunit, allowing for subsequent recruitment of mRNAs, thereby setting up a mechanism to ensure only mature subunits enter the translating pool.

On 40S subunits, Nob1 has some steric overlap with the eIF3a subunit of the translation initiation factor eIF3 (Ameismeier et al., 2018; Strunk et al., 2011). eIF3 is essential for recruiting mRNA and the ternary complex to the 40S subunit during translation (Hinnebusch and Lorsch, 2012). Furthermore, the platform region, where Nob1 is located, might also be the site of interaction with the cap-binding complex. Therefore, Nob1-bound pre-40S ribosomes would also be unable to engage in translation.

### The Rio1 kinase licenses nascent 40S ribosomes through release of Nob1 and Pno1

While rRNA cleavage is necessary for the dissociation of Nob1 from nascent ribosomes, it is not sufficient; likely because binding interactions with Pno1 keep it bound to the nascent 40S (Woolls et al., 2011). Indeed, our genetic and biochemical data demonstrate that Rio1 uses ATP hydrolysis to release both Nob1 and its binding partner Pno1 from nascent ribosomes.

Previous work has shown that Rio1 associates with late pre-40S subunits that retain only Nob1 and Pno1 (Turowski et al., 2014; Widmann et al., 2012). The data herein demonstrate that Rio1 directly binds a complex of Nob1•Pno1 in an ATP-dependent manner, and that Rio1 uses ATP hydrolysis to release both factors from the ribosome. Finally, Rio1 overexpression suppresses the dominant-negative phenotype of the Nob1-D15N mutation, and weakly binding Pno1 and Nob1 mutant proteins bypass the requirement for Rio1 *in vivo*. Together these data establish a role for Rio1 in release of Nob1 and Pno1 from nascent ribosomes, thereby regulating their entry into the translating pool in an ATPase-dependent manner. This role for Rio1 is consistent with data in the human system that show Nob1 and Pno1 re-import into the nucleolus is more strongly affected by mutations in the Rio1 active site than other assembly factors (Widmann et al., 2012).

### How does Rio1 release Nob1 and Pno1?

Rio1 is an atypical aspartate kinase. Analogous to its close relative, Rio1 is believed to use a cycle of autophosphorylation and subsequent dephosphorylation to promote its function in 40S ribosome biogenesis (Ferreira-Cerca et al., 2012; Knuppel et al., 2017). Based on our binding and release data, as well as these considerations, we speculate that ATP-bound Rio1 binds ribosome-bound Nob1•Pno1. Phosphorylation of Rio1 (and presumably release of the ADP) are then required to promote a conformational change, which leads to dissociation of the complex, with the cycle being re-set by Rio1 dephosphorylation.

### Rps26 marks matured ribosomes that passed Rio1-mediated quality control

Pno1 and Rps26 bind overlapping sites on 40S ribosomes, and as a result, Pno1-containing ribosomes lack Rps26 (Heuer et al., 2017; Strunk et al., 2011). Thus, Rps26 incorporation requires Rio1-dependent release of Pno1. In addition, our data also suggest that 20S-containing immature 40S cannot bind Rps26, consistent with structural data that indicate physical overlap (Ameismeier et al., 2018), and previous TAP-pulldowns, which indicate that Rps26 incorporation occurs after 18S rRNA maturation, as they show that ribosomes from Rps26-depleted cells contain mature 18S (Ferreira-Cerca et al., 2007; Ferretti et al., 2017). Thus, Nob1-dependent rRNA maturation is a prerequisite for both its own release, as well as Rps26 binding. Thus, we can conclude that Rps26 marks ribosomes as mature.

We have previously shown that Rps26 is not required for translation initiation (Ferretti et al., 2018; Ferretti et al., 2017), although we cannot rule out the possibility that translation initiation occurs via alternative pathways, akin to those under stress. Nonetheless, these data demonstrate that the Rps26-mark is not required to enable translation. We speculate that marking mature 40S subunits with Rps26 may render these ribosomes more stable within the cell, while Rps26-deficient ribosomes may be more likely targeted for degradation. This is consistent with the reduced number of ribosomes in Rps26-deficient yeast (Ferretti et al., 2017).

### A quality control checkpoint is established by Nob1 and Pno1, and regulated by Rio1

Bypassing Rio1 via a self-releasing Pno1 mutant allows immature 20S pre-rRNA to escape into the polysomes. Furthermore, purified ribosomes from these strains have substoichiometric Rps26 levels, resistance to high salt and high pH, and translational fidelity defects akin to Rps26-deficient cells. Thus, the data herein demonstrate a critical role for Rio1 in ensuring only fully matured ribosomes enter the translating pool. In addition, these data also show how in certain cases, Rps26-deficient ribosomes can be formed *de novo* during ribosome synthesis rather than by removing Rps26 from existing mature ribosomes.

Collectively, our data support a model (**Figure 6, top**) by which the assembly factor Pno1 and immature rRNA cooperate to block Rps26 incorporation, while the endonuclease Nob1 blocks premature mRNA recruitment. Nob1 release and Rps26 incorporation both require rRNA maturation, while Nob1 and Pno1 release require the ATP-hydrolysis activity of Rio1. Thus, we suggest that after Nob1-dependent cleavage of 20S pre-rRNA into mature 18S rRNA, Rio1 releases both Nob1 and Pno1 from nascent 40S subunits, allowing for the recruitment of Rps26 and mRNA, and the first round of translation by newly-made 40S ribosomes.

**Figure 6.**
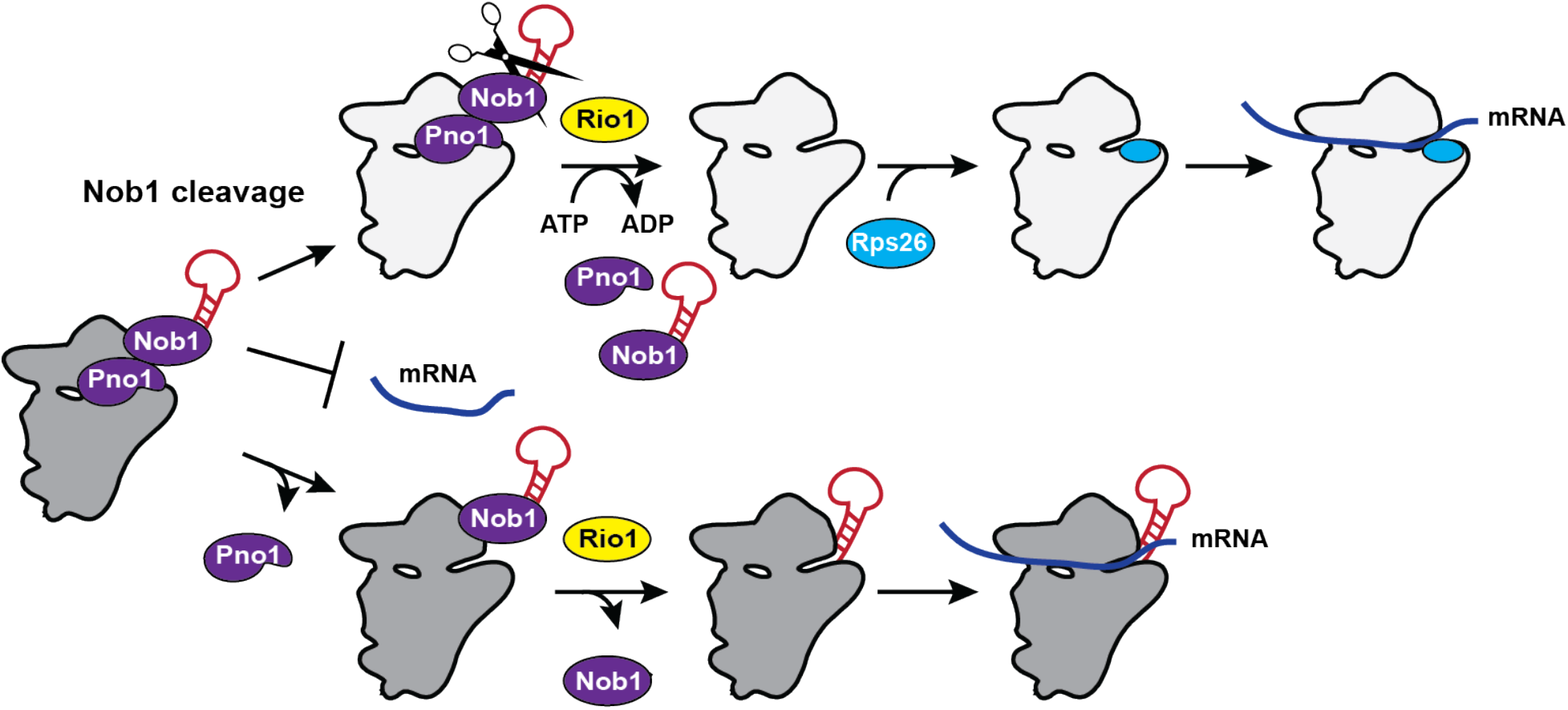
Model of Rio1’s role in 40S ribosome biogenesis. Top: Nob1 blocks mRNA recruitment. Following Nob1-mediated cleavage of 20S pre-rRNA to form the mature 18S rRNA, Rio1 binds the 40S-bound Nob1•Pno1 complex and releases both assembly factors from the nascent ribosome in an ATP-dependent manner. This allows for Rps26 incorporation and mRNA recruitment. Therefore, Rio1 regulates the final stages of 40S maturation in the cytoplasm to only release fully mature, Rps26-containing ribosomes into the translating pool. Bottom: When Rio1 is bypassed, such as in cells expressing the weak-binding Pno1-KKKF, Pno1 can dissociate even before Nob1-dependent rRNA cleavage. Because Pno1 is required for D-site cleavage (Woolls et al., 2011), this reduces the endonuclease activity of Nob1. Eventually Nob1 can be released, even when RNA cleavage has not occurred, allowing mRNA recruitment by 20S pre-rRNA-containing and Rps26-deficient 40S ribosomes.

The importance of this safeguard is demonstrated in cells expressing the self-releasing Pno1 mutant Pno1-KKKF, which bypasses Rio1’s function and rescues the lethal effect from Rio1 depletion. In these cells Pno1 can dissociate Rio1-independently and does so in a fraction of the molecules even before Nob1 matures 18S rRNA (**Figure 6, bottom**). Pno1 is required for Nob1-mediated 18S rRNA cleavage (Woolls et al., 2011). Thus, in ribosomes where Pno1 has dissociated prematurely, rRNA cleavage by Nob1 is impaired. Nonetheless, because Pno1 forms a direct interaction with Nob1 on the pre-40S ribosome (Ameismeier et al., 2018; Johnson et al., 2017; Strunk et al., 2011), and changes its RNA binding mode to strengthen Nob1’s binding affinity (Woolls et al., 2011), Pno1 dissociation will weaken Nob1 binding leading to its release from pre-40S before rRNA cleavage. As a result of Nob1 dissociation, the 20S pre-rRNA-containing pre-ribosomes can enter the translating pool, and because they remain immature, Rps26 cannot be incorporated. Thus, Nob1 and Pno1 cooperate to block premature release of immature 40S subunits into the translating pool, and Rio1 regulates the passage through this checkpoint. Whether Rio1 relies simply on the reduced affinity of Nob1 for cleaved rRNA for its temporal regulation, or actively recognizes cleaved rRNA will require further studies, including perhaps structural work.

### Why do 20S pre-rRNA-containing ribosomes not support cell growth?

Although the weakly binding Nob1-363 can bypass the requirement for Rio1 in release of rRNA into the polysomes, it does not rescue the cell viability defect from Rio1-deficient cells. Similarly, Nob1-deficient yeast accumulate ribosomes that can translate, but not support cell viability. The data herein show that in both cases these immature ribosomes retain Pno1, thus preventing the incorporation of Rps26. Therefore, while a self-releasing Nob1 mutant can dissociate Rio1-independently, Pno1 dissociation requires Nob1 and Rio1. Our data demonstrate that Rio1 recognizes the interface between Nob1 and Pno1. Thus, in the absence of Nob1, Pno1 cannot be released in a Rio1-dependent manner. It appears that Pno1 is less sensitive to the presence of Nob1 in its own binding, consistent with the recruitment of Pno1 to pre-40S prior to Nob1 (Chaker-Margot et al., 2015; Schäfer et al., 2003; Zhang et al., 2016).

### Bypassing the Rio1 checkpoint disturbs protein homeostasis and can promote cancer

The importance of the Rio1-dependent checkpoint for protein homeostasis is demonstrated by the effects from its bypass on translational fidelity and yeast biology. Depletion of Rps26, depletion of Nob1, or directly bypassing Rio1 each lead to defects in codon recognition during translation. Moreover, depletion of Rps26 or Rio1 also produce unique growth phenotypes, which are likely due to the accumulation of Rps26-deficient ribosomes, and their effects on mRNA selectivity (Ferretti et al., 2017).

Rio1 is conserved throughout all domains of life. While Rio1 is critical for ribosome assembly and cell viability in yeast, it plays an equally important role during ribosome assembly in human cells (Weinberg et al., 2017). Intriguingly, whole genome sequencing of cancer samples reveals that diverse cancers accumulate mutations in Pno1 that are either directly adjacent to Pno1-KKKF, or similarly contact either the rRNA, Nob1, or ribosomal proteins (**Figure S6**, TCGA Research Network: https://www.cancer.gov/tcga). Thus, while it remains unclear whether these mutations play any role in promoting cancer progression, like Pno1-KKKF, the cancer-associated Pno1 mutants are expected to bypass Rio1, leading to the release of immature, Rps26-deficient ribosomes into the translating pool, as we have shown in yeast cells. Importantly, Rps26 haploinsufficiency results in a predisposition to the development of specific forms of cancer (Armistead and Triggs-Raine, 2014; Sulima et al., 2017; Vlachos, 2017), and furthermore, ribosomes from cancer cells often lose the stoichiometry of ribosomal proteins, including Rps26, which is correlated with poor patient outcomes (Ajore et al., 2017; Guimaraes and Zavolan, 2016; Kulkarni et al., 2017; Vlachos, 2017). Together, these data argue for a role for Rps26-deficient cells in cancer progression. Furthermore, these data also demonstrate that cancer cells have found diverse means to effect loss of ribosomal protein stoichiometry, including changes in RP gene expression (Ajore et al., 2017; Guimaraes and Zavolan, 2016; Kulkarni et al., 2017; Vlachos, 2017), downregulation of the assembly factor Ltv1 leading to substoichiometric, Rps3, Rps10 and Asc1/RACK1 (Collins et al., 2018), and bypass of Rio1, leading to Rps26-deficient ribosomes. This observation is consistent with a general role for many ribosomal proteins as “tumor suppressors” (Amsterdam et al., 2004).

## Materials and Methods

### Yeast strains and cloning

*Saccharomyces cerevisiae* strains used in this study were obtained from the GE Dharmacon Yeast Knockout Collection or were made using PCR-based recombination (Longtine et al., 1998). Strain identity was confirmed by PCR and Western blotting when antibodies were available. Mutations in plasmids were made by site-directed mutagenesis and confirmed by sequencing. Rio1 was cloned into pSV272, for expression as a TEV-cleavable His_6_-MBP fusion protein. Plasmids were propagated in XL1 Blue competent cells. Yeast strains and plasmids used in this study are listed in Tables S1 and S2, respectively.

### Protein expression and purification

Pno1, MBP-Pno1, Nob1, and MBP-Nob1 were purified as previously described (Campbell and Karbstein, 2011; Lamanna and Karbstein, 2009; Woolls et al., 2011). Truncated Nob1-363 was purified using the same protocol as the wild-type protein.

To express and purify Rio1, Rosetta DE3 competent cells transformed with a plasmid encoding His-MBP-tagged Rio1 were grown to mid-log phase at 37°C in LB media supplemented with the appropriate antibiotics. Rio1 expression was induced by addition of 1mM isopropyl β-D-thiogalactoside (IPTG) and cells grown for another 5h at 30°C. Cells were lysed by sonication in Ni-NTA lysis buffer supplemented with 0.5 mM phenylmethylsulfonyl fluoride (PMSF) and 1 mM benzamidine. The cleared lysate was purified over Ni-NTA affinity resin according to the manufacturer’s recommendation (Qiagen). Eluted proteins were pooled and dialyzed overnight at 4°C into 50 mM Na_2_HPO_4_ pH 8.0, 150 mM NaCl, 1 mM DTT. Protein was applied to a MonoQ column in the same buffer and eluted with a linear gradient of 150 mM to 600 mM NaCl over 12 column volumes. The protein was pooled and concentrated for further purification on a Superdex200 size-exclusion column equilibrated with (50 mM Hepes pH 8.0, 200 mM NaCl, 1 mM DTT, 1 mM TCEP). Protein concentration was determined by absorption at 280 nm using an extinction coefficient of 106,120 M^−1^cm^−1^.

Untagged Rio1 was purified as described above, except that 0.76 μg/mL TEV protease was added during dialysis. Protein concentration was determined by absorption at 280 nm using an extinction coefficient of 36,790 M^−1^cm^−1^.

### Sucrose density gradient analysis

Sucrose gradient fractionation of whole cell lysates followed by Northern blot analysis were performed as described previously (Strunk et al., 2012). Briefly, cells were grown to mid log phase in the appropriate media (indicated in the respective figure legends), harvested in 0.1 mg/mL cycloheximide, washed, and lysed in gradient buffer (20 mM Hepes, pH 7.4, 5 mM MgCl_2_, 100 mM KCl, and 2 mM DTT) with 0.1 mg/mL cycloheximide, complete protease inhibitor cocktail (Roche), 1 mM benzamidine, and 1 mM PMSF. Cleared lysate was applied to 10-50% sucrose gradients and centrifuged in an SW41Ti rotor for 2h at 40,000 RPM and then fractionated. The percent of 20S pre-rRNA in the polysomes was calculated by dividing the amount of 20S pre-rRNA in the polysome fractions (fractions 8-13) by the total amount of 20S pre-rRNA in all fractions (fractions 2-13).

### Protein binding assays

7 μM of MBP-tagged protein (MBP-Rio1, MBP-Pno1, and MBP-Nob1) was mixed with 20 μM untagged protein (Rio1, Nob1, or Pno1) in binding buffer (50 mM Hepes pH 7.5, 200 mM NaCl, and 5 mM MgCl_2_). 2 mM ATP or AMPPNP was added where indicated. Proteins were pre-incubated at 4°C for 30 min before addition of 100 uL equilibrated amylose resin (New England BioLabs). The mixture was incubated for 1h at 4°C, the flow-through was collected, the resin was washed with binding buffer supplemented with 0.8 mM ATP or AMPPNP where indicated, and proteins were eluted with binding buffer supplemented with 50 mM maltose.

### RNA binding assay

RNA binding assays were performed as previously described (Lamanna and Karbstein, 2009). Briefly, ^32^P-ATP-labeled H44-A2, H44-D, or D-A2 RNAs, named after the sequence on structural elements that mark their start and end points, were prepared by transcription in the presence of α-ATP, gel-purified and eluted via electroelution. These RNAs have been validated to fold into well-defined structures relevant to ribosome assembly (Lamanna and Karbstein, 2009, 2011). RNAs were then precipitated and resuspended in water. RNAs were folded by heating for 10 min at 65 °C in the presence of 40 mM HEPES, pH 7.6, 100 mM KCl, and 2 mM MgCl_2_. Trace amounts of radiolabeled RNA were incubated with varying concentrations of Nob1 (or Nob1•Pno1 or Nob1•Pno1•Rio1) in 40 mM HEPES, pH 7.6, 50 mM KCl and 10 mM MgCl2 for 10 min at 30°C. Samples were loaded directly onto a running 6% acrylamide/THEM native gel to separate protein-bound from unbound RNAs. After drying the gel, phosphorimager analysis was used to quantify the gel. Bound RNA was plotted against protein concentration and fit with a single binding isotherm to obtain apparent binding constants using KaleidaGraph version 4.5.4 from Synergy Software.

### Release assay

Pre-ribosomes from Rio1-depeleted cells were purified from Gal::Pno1; Gal::Rio1 cells transformed with a plasmid encoding Pno1-TAP and grown in YPD-medium for 16 h essentially as described before (Ghalei et al., 2017). 40 nM of pre-40S ribosomes were incubated with 2 μM purified, recombinant Rio1 in 50 μL buffer (50 mM Tris-HCl, pH 7.5, 100 mM NaCl, 10 mM MgCl_2_, 0.075% NP-40, 0.5 mM EDTA, and 2 mM DTT). ATP, AMPPNP, or ADP were added to a final concentration of 1 mM. The samples were then incubated at room temperature for 10 min, placed on 400 μL of a 20% sucrose cushion, and centrifuged for 2 hr at 400,000 g in a TLA 100.1 rotor. The supernatant was TCA-DOC precipitated and the pellets were resuspended in SDS-loading dye. Supernatants (released factors) and pellets (bound factors) were analyzed by SDS-PAGE followed by Western blotting.

### Ribosome purification

40S ribosomes were purified from Gal::Pno1 cells expressing wt Pno1 or Pno1-KKKF as described previously (Acker et al., 2007). Briefly, cells were grown to OD_600_ 1.5-1.7 and flash-frozen in ribosome buffer (20 mM Hepes pH 7.4, 100 mM KOAc, 2.5 mM Mg(OAc)_2_) supplemented with 1 mg/mL heparin, complete protease inhibitor cocktail (Roche), 1 mM benzamidine, and 1 mM PMSF. 3 mL cleared lysate was layered over 0.5 mL of sucrose cushion (ribosome buffer, 500 mM KCl, 1M sucrose, 2 mM DTT) and centrifuged in a TLA 110 rotor at 70,000 rpm for 65 min. The pellets were resuspended in high-salt buffer (ribosome buffer, 500 mM KCl, 1 mg/mL heparin, 2 mM DTT) and again layered over a 0.5 mL sucrose cushion and centrifuged at 100,0000 rpm for 70 min. The pellets were resuspended in subunit separation buffer (50 mM Hepes pH 7.4, 500 mM KCl, 2 mM MgCl_2_, 2 mM DTT). Once suspended, puromycin was added to a concentration of 1 mM, the sample was incubated on ice for 15 min, incubated at 37°C for 10 min, loaded onto a 5-20% sucrose gradient, and centrifuged in an SW32 rotor for 6h at 32,000 rpm. The 40S ribosome-containing fractions were pooled and analyzed by SDS-PAGE followed by Western blotting.

### Quantitative growth assays

Stress-tolerance tests were performed as previously described (Ferretti et al., 2017). In brief, Gal::Pno1 cells transformed with a plasmid encoding Pno1 or Pno1-KKKF were grown to mid-log phase in glucose dropout media and then inoculated into stress media (or control cultures) at OD 0.05 to test stress tolerance. The stress media was either YPD + 1 M NaCl (high salt) or YPD + 100 mM TAPS, pH 8.2 (high pH). YPD was used as the control media for the high salt conditions, while YPD + 100 mM TAPS, pH 7.0 was used as the control media for the high pH conditions. Cells were grown at 30°C while shaking and the doubling times were measured in a Synergy 2 multi-mode microplate reader (BioTek).

### Dual-luciferase reporter assay

Gal::Pno1 cells grown in glucose media were supplemented with plasmids encoding either wt Pno1 or Pno1-KKKF. Gal::Nob1 cells grown in glucose media were supplemented with either wt Nob1 or an empty vector. ΔRps26B; Gal::Rps26A cells supplemented with plasmids encoding Rps26A under a constitutively active TEF promoter or a doxycycline-repressible TET promoter. Cells were harvested in mid-log phase and reporter assays carried out essentially as described before (Ghalei et al., 2017). Cells were lysed, and luciferase activity was measured with the Promega Dual-Luciferase Reporter Assay System on a PerkinElmer EnVision 2104 Multilabel Reader according to the manufacturer’s protocol with assay volumes scaled down to 15%. For each sample, firefly luciferase activity was normalized against renilla activity, subsequently, values observed for mutant Pno1 were normalized against those for wt Pno1. For Rps26 cells, values observed for depleted Rps26 were normalized against those for wt Rps26. For Nob1 cells, values observed for depleted Nob1 were normalized against those for wt Nob1.

### Antibodies

Antibodies against recombinant Nob1, Pno1, Rps10, and Rps26 were raised in rabbits by Josman LLP or New England Peptide, and tested against purified recombinant proteins and yeast lysates.

### Quantification and Statistical Analysis

Quantification of Northern and Western blots was performed using Quantity One 1-D Analysis Software version 4.1.2 (Basic) and Image Lab version 5.2.1, respectively, from Bio-Rad Laboratories, Inc. Statistical analysis of the dual luciferase translation fidelity assay was performed using GraphPad Prism version 6.02, GraphPad Software, La Jolla, California USA, www.graphpad.com. Statistical analyses of Northern blots and growth assays were performed using the programming language R in Rstudio, version 3.2.3 (R Core Team (2015). R: A language and environment for statistical computing. R Foundation for Statistical Computing, Vienna, Austria. URL https://www.R-project.org/). Samples grown and analyzed on the same day were considered paired replicates and significance was calculated using a paired, two-tailed t-test. Otherwise, unpaired, two-tailed t-test was used as indicated in the figure legends.

## Acknowledgements

This work was supported by National Institute of Health grant R01-GM086451 and HHMI Faculty Scholar Grant 55108536 (to K.K.). We thank A. Lamanna for the gift of recombinant Nob1-363, and G. Dieci, A. Link, L. Valášek, and J. Warner for gifting us the anti-Rps8, anti-Asc1, anti-Rps0, and anti-Rps2 antibodies, respectively. The dual luciferase plasmids were kindly provided by J. Lorsch (start site recognition), D. Bedwell (stop codon read-through and miscoding), and J. Dinman (frameshifting). We thank T. Mueller and members of the Karbstein lab for discussion and comments on the manuscript.

## Author Contributions

M.D.P., J.C.C., B.K., H.G., and K.K. designed and performed the experiments. M.P. and K.K.

## Supplemental Information

**Figure S1.**
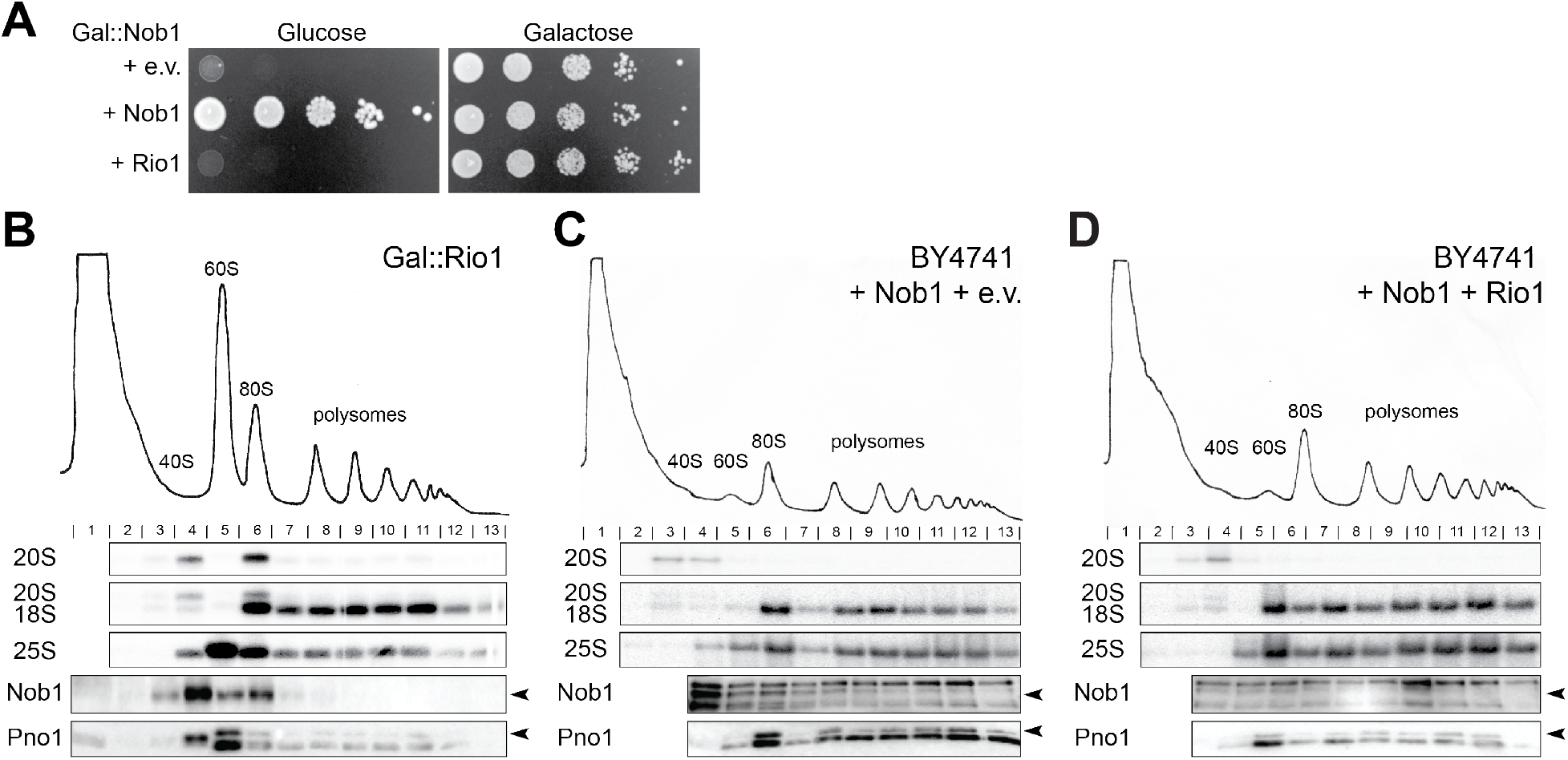
Rio1 depletion and the Nob1-D15N mutation result in a similar phenotype. Related to Figure 1 and 3. **A)** Overexpression of Rio1 does not rescue Nob1 depletion. Growth of cells containing Nob1 under a Gal promoter and expressing either Nob1 or Rio1 from a plasmid under a copper-inducible (Cup1) promoter or an empty vector were compared by 10-fold serial dilutions on glucose or galactose dropout plates with 100 μM CuSO_4_. **B)** 10-50% sucrose gradient from cell lysate of Gal:Rio1 cells depleted of Rio1 by growth in YPD for 16 hr. Northern blots of 20S, 18S, and 25S rRNA and Western blots probing for Nob1 and Pno1 are shown below the absorbance profile at 254 nm. 19% (+/− 1%) of 20S accumulated in the polysomes (fractions 8-13). **C)** Sucrose gradient from wild type cells transformed with an empty vector and overexpressing wild type Nob1 under a Gal promoter grown in galactose with 100 μM CuSO4 for 16 hr. **D)** Sucrose gradient from wild type cells overexpressing Nob1-D15N under a Gal promoter and Rio1 under a copper-inducible (Cup1) promoter grown in galactose with 100 μM CuSO_4_ for 16 hr. Arrowheads note the bands corresponding to Nob1 and Pno1.

**Figure S2.**
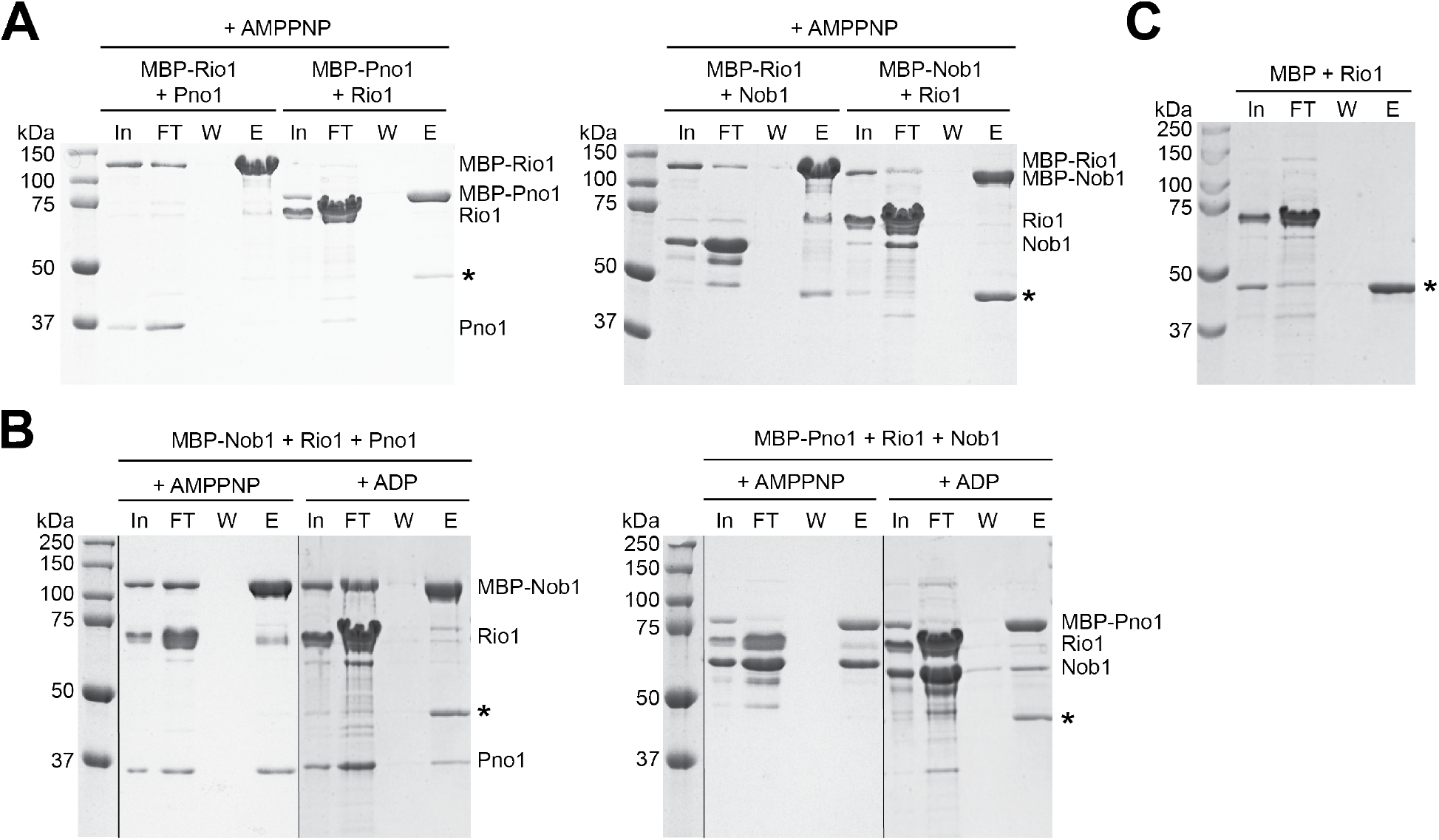
Rio1 does not bind Nob1 or Pno1 individually. Related to Figure 4. **A)** Rio1 does not bind Nob1 or Pno1 individually. Shown are coomassie-stained SDS-PAGE gels of protein binding assays of purified, recombinant MBP-Rio1, Nob1, MBP-Nob1, Rio1, Pno1, and MBP-Pno1 in the presence of AMPPNP. **B)** Coomassie-stained SDS-PAGE gels of protein binding assays on amylose beads of purified, recombinant MBP-Nob1, Rio1, Pno1, MBP-Pno1, and Nob1 in the presence of AMPPNP or ADP. In, input; FT, flow-through; W, final wash; E, elution. The order of the samples was edited for clarity. **C)** Rio1 does not bind MBP. Shown is a coomassie-stained SDS-PAGE gel of a protein binding assay of purified, recombinant MBP and Rio1. Nob1 and Pno1 also do not bind MBP alone (Woolls et al., 2011). *MBP

**Figure S3.**
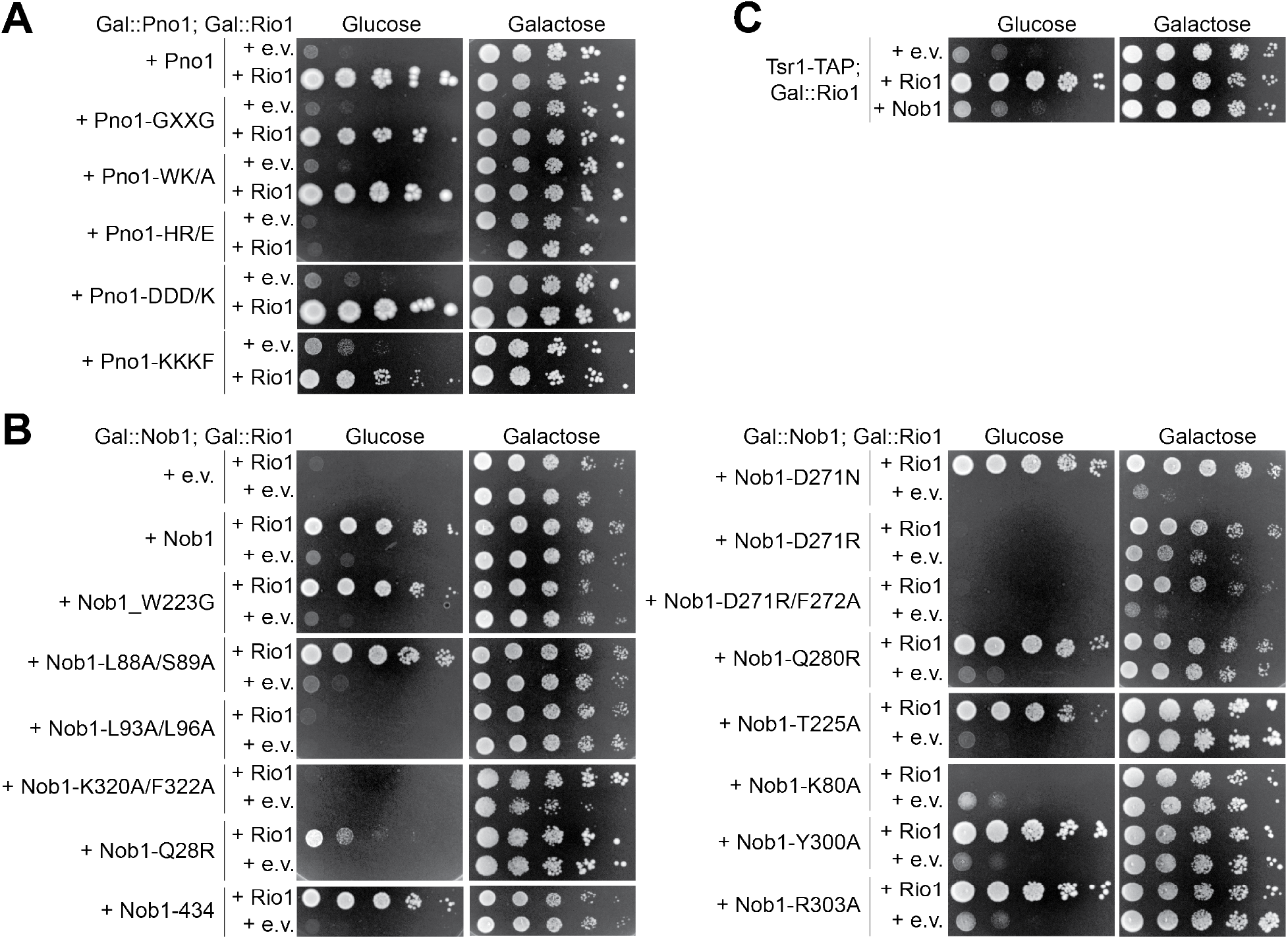
Rescue of the Rio1 depletion phenotype is specific to Pno1-KKKF. Related to Figure 4. **A)** Growth of cells expressing wild-type Pno1 or Pno1 mutants with and without Rio1 were compared by 10-fold serial dilutions on glucose and galactose dropout plates. Pno1-GXXG (N111G/S112K/W113D/T114G), Pno1-WK/A (W113A/K115A), Pno1-HR/E (H104E/R105E), Pno1-DDD/K (D167K/D169K/D170K). **B)** Growth of cells expressing wild-type Nob1 or Nob1 mutants with or without Rio1 were compared by 10-fold serial dilutions on glucose and galactose dropout plates. **C)** Growth of cells containing endogenous Rio1 under a Gal promoter expressing either wild-type Nob1 or Rio1 under a copper-inducible (Cup1) promoter or an empty vector were compared by 10-fold serial dilutions on glucose or galactose dropout plates with 100 μM CuSO_4_.

**Figure S4.**
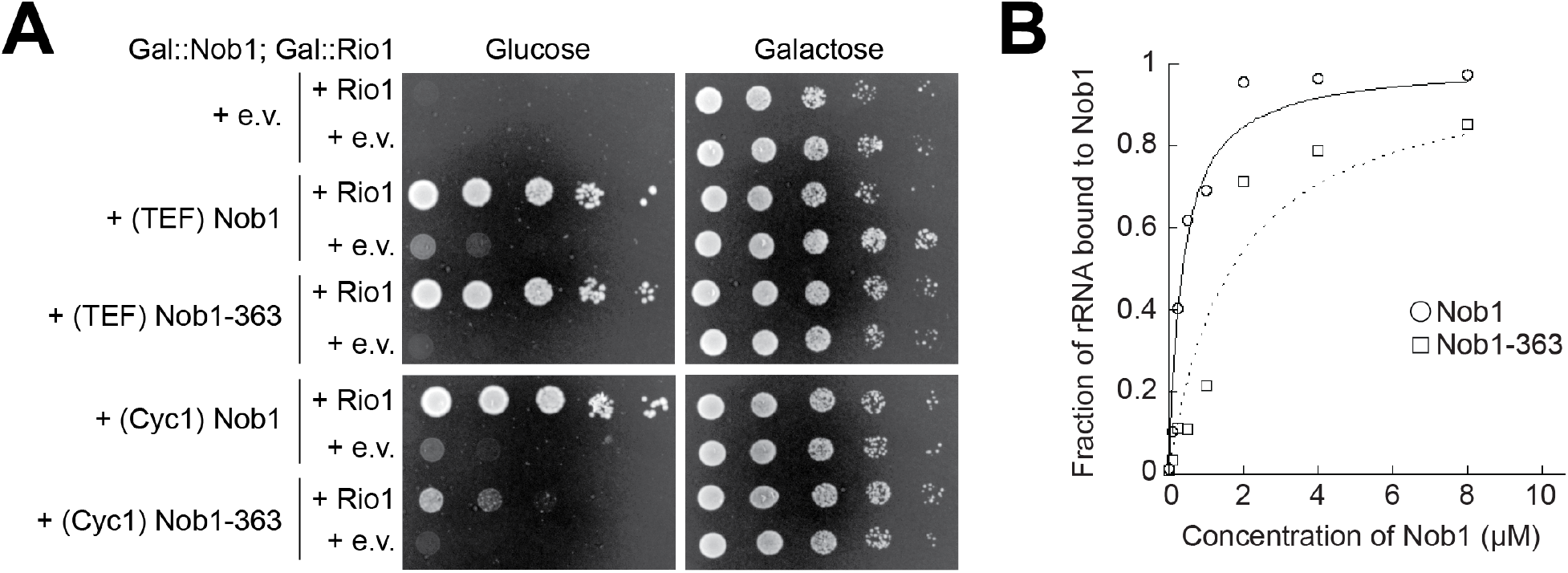
Truncated Nob1-363 weakly binds RNA. Related to Figure 4. Growth of cells expressing wild-type Nob1 or Nob1 mutants under a TEF or Cyc1 promoter, as indicated, with or without Rio1 were compared by 10-fold serial dilutions on glucose and galactose dropout plates. TEF promoters produce higher protein levels (Mumberg et al., 1995). B) RNA binding assay with *in vitro* transcribed H44-A2 RNA (20S pre-rRNA mimic) and recombinant Nob1 or Nob1-363. Three independent experiments yielded values of *K*_d_ = 0.37 +/− 0.22 μM for Nob1 binding H44-A2 (white circles) and *K*_d_ = 1.43 +/− 0.22 μM for Nob1-363 binding H44-A2 (white squares).

**Figure S5.**
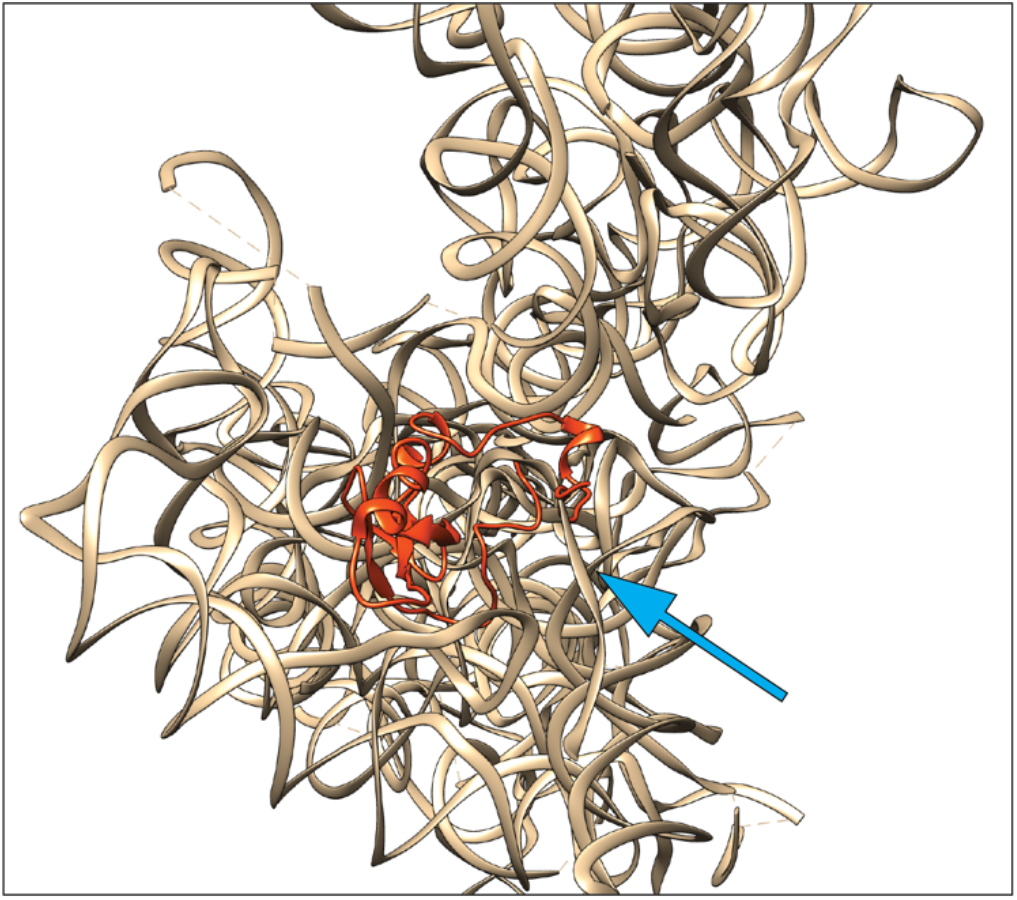
Rps26 cannot bind 40S ribosomes containing pre-18S rRNA. Related to Figure 5. Rps26 (from mature human 40S, in red) overlaps the 3’-end of pre-18S rRNA, marked by the blue arrow (from human pre-40S, tan). Image was obtained by overlaying 18S rRNAs from PDBID 6G5H and 6G18 (mature human 40S and human pre-40S state C, respectively, (Ameismeier et al., 2018)) using the matchmaker tool in Chimera. For simplicity all proteins other than Rps26 are omitted.

**Figure S6.**
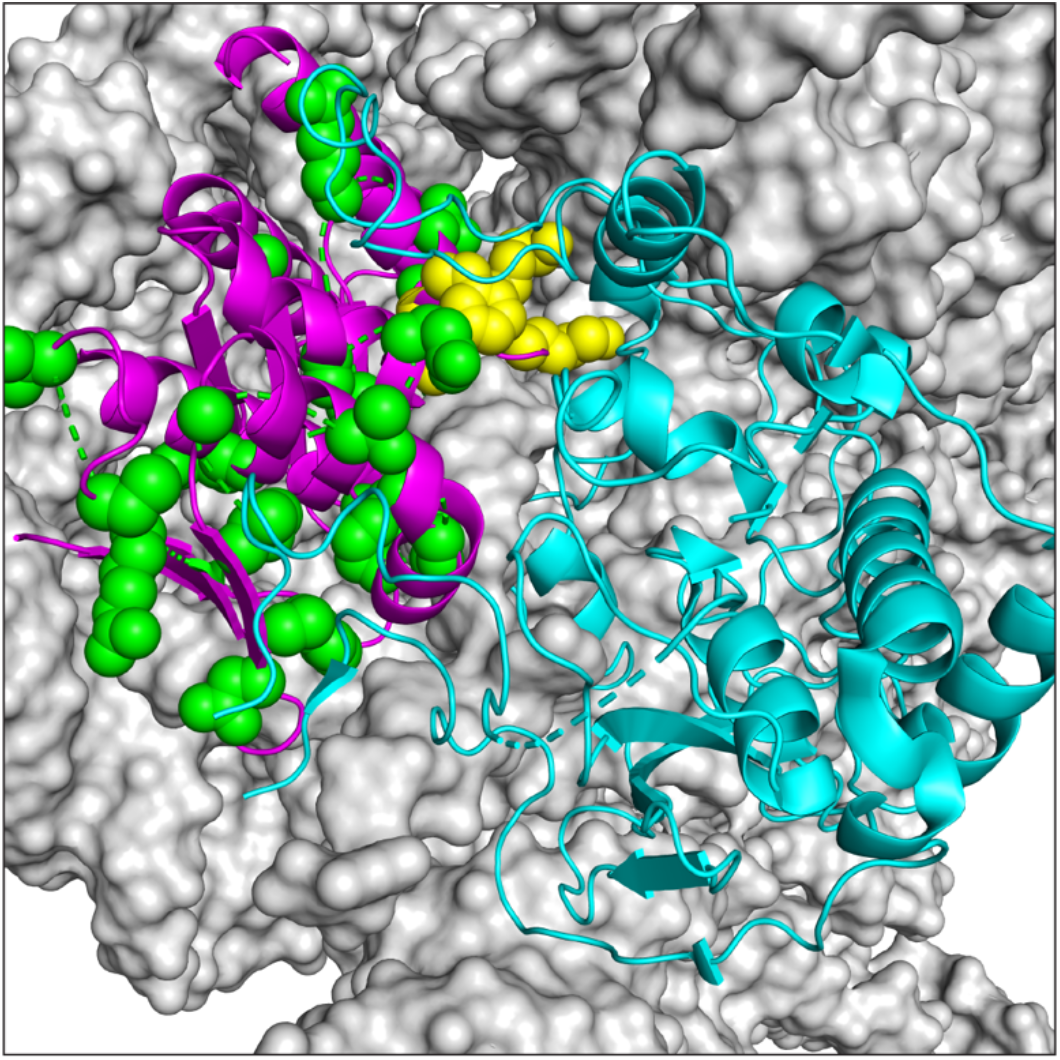
Pno1 mutations discovered in cancer genomes. Related to the Discussion. Mutations in Pno1 that accumulate in diverse cancers (green space fill, from the TCGA Research Network: https://www.cancer.gov/tcga) are directly adjacent to Pno1-KKKF (yellow space fill) or similarly contact either the rRNA, Nob1, or ribosomal proteins accumulate in diverse cancers. Pre-mature 18S rRNA (from human pre-40S, surface view in grey) bound by Nob1 (cyan) and Pno1 (magenta). Image was obtained from PDBID 6G18 (human pre-40S state C, (Ameismeier et al., 2018)). For simplicity, all proteins other than Nob1 and Pno1 are omitted.

**Table S1:** Yeast strains used in this work

| **Strain** | **Description** | **Background** | **Genotype** | **Reference** |
| --- | --- | --- | --- | --- |
| YKK200 | WT | BY4741 | *MATα his3Δ1 leu2Δ0 met15Δ0 ura3Δ0* | GE Dharmacon |
| YKK650 | Gal::Nob1 | BY4741 | *MATα KanMX6::pGAL1-Nob1 his3Δ1 leu2Δ0 met15Δ0 ura3Δ0* | This work |
| YKK487 | Gal::Pno1 | BY4741 | *MATα KanMX6::pGAL1-Pno1 his3Δ1 leu2Δ0 met15Δ0 ura3Δ0* | (Johnson et al., 2017) |
| YKK281 | Gal::Rio1<br>Tsr1-TAP | BY4741 | *MATα KanMX6::pGAL1-Rio1 his3::Tsr1-TAP leu2Δ0 met15Δ0 ura3Δ0* | This work |
| YKK1134 | Gal::Nob1<br>Gal::Rio1 | BY4741 | *MATα KanMX6::pGAL1-Nob1 NatMX6::pGal1-Rio1 his3Δ1 leu2Δ0 met15Δ0 ura3Δ0* | This work |
| YKK988 | Gal::Pno1<br>Gal::Rio1 | BY4741 | *MATα KanMX6::pGAL1-Pno1 NatMX6::pGAL1-Rio1 his3Δ1 leu2Δ0 met15Δ0 ura3Δ0* | This work |
| YKK491 | Gal::Rps26 | BY4741 | *MATα NatMX6::pGAL1- Rps26A Rps26B::KanMX6 his3Δ1 leu2Δ0 met15Δ0 ura3Δ0* | (Ferretti et al., 2017) |

**Table S2:** Plasmids used in this work

| **Plasmid** | **Description** | **Backbone** | **Reference** |
| --- | --- | --- | --- |
| pKK3464 | Gal::Nob1 | pRS426 | This work |
| pKK3723 | Gal::Nob1-D15N | pRS426 | This work |
| pKK30014 | CUP1::Nob1 | pRS426 | This work |
| pKK3387 | TEF::Nob1 | pRS413 | This work |
| pKK30177 | TEF::Nob1-W223G | pRS413 | This work |
| pKK378 | TEF::Nob1 | pRS416 | This work |
| pKK30399 | TEF::Nob1-L88A/S89A | pRS416 | This work |
| pKK30400 | TEF::Nob1-L93A/L96A | pRS416 | This work |
| pKK3056 | TEF::Nob1-K320A/F322A | pRS416 | This work |
| pKK30401 | TEF::Nob1-D271N | pRS416 | This work |
| pKK30402 | TEF::Nob1-D271R | pRS416 | This work |
| pKK30403 | TEF::Nob1-D271R/F272A | pRS416 | This work |
| pKK30404 | TEF::Nob1-Q280R | pRS416 | This work |
| pKK3243 | TEF::Nob1-K80A | pRS416 | This work |
| pKK3246 | TEF::Nob1-R303A | pRS416 | This work |
| pKK3245 | TEF::Nob1-Y300A | pRS416 | This work |
| pKK3211 | TEF::Nob1-363 | pRS416 | This work |
| pKK377 | CYC1::Nob1 | pRS415 | This work |
| pKK396 | CYC1::Nob1-363 | pRS415 | This work |
| pKK397 | CYC1::Nob1-434 | pRS415 | This work |
| pKK381 | CYC1::Nob1-T225A | pRS415 | This work |
| pKK3124 | CYC1::Nob1-Q28R | pRS415 | This work |
| pKK30156 | TEF::Rio1 | pRS415 | This work |
| pKK3280 | TEF::Rio1 | pRS416 | This work |
| pKK3999 | CUP1::Rio1 | pRS426 | This work |
| pKK30028 | CUP1::Rio1 | pRS423 | This work |
| pKK30073 | CUP1::Rio1-D261A | pRS423 | This work |
| pKK30074 | CUP1::Rio1-K86A | pRS423 | This work |
| pKK3270 | TEF::Pno1 | pRS416 | (Woolls et al., 2011) |
| pKK3823 | TEF::Pno1-KKKF | pRS416 | (Johnson et al., 2017) |
| pKK3271 | TEF::Pno1-GxxG | pRS416 | (Woolls et al., 2011) |
| pKK3272 | TEF::Pno1-WK/A | pRS416 | This work |
| pKK3273 | TEF::Pno1-HR/E | pRS416 | (Woolls et al., 2011) |
| pKK3274 | TEF::Pno1-DDD/K | pRS416 | (Woolls et al., 2011) |
| pKK30157 | TEF::TAP-Pno1 | pRS413 | This work |
| pKK3792 | TET::Rps26A | pCM189 | (Ferretti et al., 2017) |
| pKK3558 | TEF::Rps26A | pRS416 | (Ferretti et al., 2017) |
| pKK30059 | ADH1/GPD-Start-site<br>(Renilla(AUG)/Firefly(UUG)) | pRS415 | (Cheung et al., 2007) |
| pKK3920 | ADH1-0 Frameshift Control | pRS415 | (Harger and Dinman, 2003) |
| pKK3921 | ADH1-liter<sub>A</sub> (-1) Frameshift | pRS415 | (Harger and Dinman, 2003) |
| pKK3922 | ADH1-Ty1 (+1) Frameshift | pRS415 | (Harger and Dinman, 2003) |
| pKK3923 | TEF-Read-through | pRS415 | (Keeling et al., 2004) |
| pKK3924 | TEF-Stop | pRS415 | (Keeling et al., 2004) |
| pKK3925 | TEF-Misincorporation<br>(Firefly_H245R) | pRS415 | (Salas-Marco and Bedwell, 2005) |
| pKK1181 | MBP-Rio1 | pSV272 | This work |
| pKK187 | MBP-Nob1 | pSV272 | (Lamanna and Karbstein, 2009) |
| pKK1051 | MBP-Nob1-363 | pSV272 | This work |
| pKK191 | MBP-Pno1 | pSV272 | (Woolls et al., 2011) |

